# Spatial differences in stoichiometry of EGR phosphatase and Microtubule-Associated Stress Protein 1 control root meristem activity during drought stress

**DOI:** 10.1101/2021.10.06.463370

**Authors:** Toshisangba Longkumer, Chih-Yun Chen, Marco Biancucci, Govinal Badiger Bhaskara, Paul E. Verslues

## Abstract

During moderate severity drought and low water potential (ψ_w_) stress, poorly understood signaling mechanisms restrict both meristem cell division and subsequent cell expansion. We found that the Clade E Growth-Regulating 2 (*EGR2*) protein phosphatase and *Microtubule Associated Stress Protein 1* (*MASP1*) differed in their stoichiometry of expression across the root meristem and had opposing effects on root meristem activity at low ψ_w_. Ectopic MASP1 or EGR expression increased or decreased, respectively, root meristem size and root elongation during low ψ_w_ stress. This, along with the ability of phosphomimic MASP1 to overcome EGR suppression of root meristem size and observation that ectopic EGR expression had no effect on unstressed plants, indicated that during low ψ_w_ EGR activation and attenuation of MASP1 phosphorylation in their overlapping zone of expression determines root meristem size and activity. Ectopic EGR expression also decreased root cell size at low ψ_w_. Conversely, both the *egr1-1egr2-1* and *egr1-1egr2-1masp1-1* mutants had similarly increased root cell size; but, only *egr1-1egr2-1* had increased cell division. These observations demonstrated that EGRs affect meristem activity via MASP1 but affect cell expansion via other mechanisms. Interestingly, EGR2 was highly expressed in the root cortex, a cell type important for growth regulation and environmental response.

**One Sentence Summary:** Spatial differences in EGR-MASP1 expression and control of MASP1 phosphorylation adjust root meristem activity to regulate growth during drought stress.

The author responsible for distribution of materials integral to the findings presented in this article in accordance with the policy described in the Instructions for Authors (www.plantcell.org) is: Paul E. Verslues.

---

Even slight or moderate reduction of water availability (quantified as a reduction in water potential, ψ_w_) leads plants to actively restrict their growth. This can be an adaptive mechanism to conserve soil water during drought periods. However, such negative regulation of growth can also restrict plant biomass accumulation more than is needed or desired in agricultural settings (Skirycz and Inzé, 2010; Verslues, 2017). A better understanding of the mechanisms that regulate plant growth in response to altered water status could allow growth inhibition to be circumvented when desirable (Skirycz and Inzé, 2010; Claeys et al., 2014).

Root growth is both a key factor in drought resistance and a model for developmental study because of the relatively simple organization of the root tip. At the root tip, the quiescent center (QC) and immediately adjacent cells act as a stem cell niche that produces cells to generate both the root meristem and root cap (Heidstra and Sabatini, 2014). Just behind the QC and adjacent cells is the meristematic region which supplies the bulk of new cells during root elongation. Cells in the meristem divide rapidly until they are displaced from the root tip, lose their cell division activity, and begin rapid expansion. The rate of root elongation is determined by both cell expansion and the amount of cell production by the meristem. Meristem cell production is in turn determined by the rate of cell division (cell cycle time) as well as the duration of time over which cell division continues as cells are displaced from the root tip. A longer duration of cell division increases the size of the root meristem (Beemster and Baskin, 1998).

While the root tip can be anatomically divided into distinct zones of cell division and cell expansion, the size of each zone may be controlled by underlying gradients of key regulatory factors (Barlow, 1976). Gradients in hormone content and signaling, particularly auxin, brassinosteroid and cytokinin, as well as reactive oxygen are important regulators of meristem size and function (Chaiwanon and Wang, 2015; Vragovic et al., 2015; Di Mambro et al., 2017; Salvi et al., 2020; Yamada et al., 2020). It is also known that hormone-related regulatory proteins and transcription factors have gradients of expression across the root meristem or are specifically expressed in the QC and surrounding stem cell area (Heidstra and Sabatini, 2014; Salvi et al., 2020; Yamada et al., 2020). Many additional genes have gradients of increasing or decreasing expression across the root meristem and some of these lesser characterized genes also affect root growth (Wendrich et al., 2017; Dubois et al., 2018; Slovak et al., 2020). It has been proposed that the balance between opposing gradients of pro-cell division genes, which are most highly expressed in the proximal meristem region close to the QC, versus pro-cell expansion genes, which are highly expressed at the distal end of the meristem and in the expansion zone, determines when cells stop dividing and begin rapid expansion (Salvi et al., 2020). Such opposing gradients of expression suggests that, at different positions within the root meristem, the stoichiometry (ratio) of the proteins will vary. Such differences in protein stoichiometry may impart differing outcomes in terms of allowing continued cell division or inhibiting cell division and promoting cell expansion (Wendrich et al., 2017). While appealing, this idea remains largely unproven as the function of most gradient-expressed genes, and whether they produce different levels of protein at different positions in the root meristem, is unclear. It is also unclear how proteins having spatial differences in stoichiometry across the meristem can affect each other’s function at the molecular level.

Low ψ_w_ during drought stress leads to both decreased cell production and decreased cell expansion (Skirycz and Inzé, 2010; Skirycz et al., 2011). Which of these is the predominant cause of reduced growth can vary between different plants and between different types of abiotic stress. In Arabidopsis roots, low ψ_w_ (−1.28 MPa), imposed using PEG-infused agar plates, led to a nearly 50% decrease in root cell production rate but only minor reduction of cell expansion (van der Weele et al., 2000). Growth kinematic analysis has also shown that drought stress decreases cell production in the root meristem (Sacks et al., 1997; Voothuluru et al., 2020). These observations suggest that stress signaling can interact with cell cycle and developmental regulation to control meristem activity. Consistent with this idea, several types of abiotic stress elicit root cell type-specific responses (Dinneny et al., 2008; Iyer-Pascuzzi et al., 2011) and development regulators can also influence stress response genes (Moubayidin et al., 2013). Despite these indications, relatively little is known of how stress signaling impinges upon developmental mechanisms to control meristem activity and regulate growth (Shimotohno and Scheres, 2019).

Previous work in our laboratory found that Clade E Growth Regulating (EGR) type 2C protein phosphatasse (PP2Cs) act as negative regulators of growth during low ψ_w_ (Bhaskara et al., 2017). Mutants of *EGR1*, *EGR2*, and *EGR3* all had higher than wild type growth (quantified based on fresh weight, dry weight and primary root length increase) and an *egr1-1egr2-1* double mutant had a stronger effect, consistent with EGR1 and EGR2 having redundant function in growth regulation during low ψ_w_. This increased growth was observed in both polyethylene glycol (PEG)-agar plate assays and in controlled soil drying experiments where plants were exposed to moderate severity water limitation. Phosphoproteomic analysis of *egr1-1egr2-1* indicated that EGRs primarily affect phosphorylation of plasma membrane, trafficking and cytoskeleton-associated proteins. One of the EGR-regulated proteins identified by phosphoproteomics was a protein of unknown function which we designated as Microtubule Associated Stress Protein 1 (MASP1). The EGR phosphatases interacted with MASP1 (in Bi-molecular Fluorescence Complementation and co-immunoprecipitation assays) and we could validate (using Phostag gel analysis) that during low ψ_w_ stress MASP1 was more phosphorylated in *egr1-1egr2-1* but less phosphorylated in transgenic plants ectopically expressing EGR1 (*35S:EGR1*). MASP1 promoted growth during low ψ_w_ in both PEG-agar plate and soil drying assays. This growth promotion activity was dependent upon MASP1 serine 670 phosphorylation (Bhaskara et al., 2017). Consistent with EGR regulation of MASP1 phosphorylation, genetic analysis indicated that MASP1 acted downstream of EGRs; although, the phenotype of the *egr1-1egr2-1masp1-1* triple mutant was somewhat intermediate between that of *egr1-1egr2-1* and *masp1-1*. MASP1 bound microtubules *in vitro*. MASP1 and EGRs had converse effects on microtubule stability *in planta* which correlated with MASP1 S670 phosphorylation status and with growth at low ψ_w_: more stable microtubules were associated with phosphorylated MASP1 and enhanced growth maintenance at low ψ_w_. Other than this, nothing is known about MASP1 cellular function. The only MASP1-related protein which has been studied is Auxin-Induced in Root Cultures 9 (AIR9), which binds microtubules and localizes at the preprophase band during cell division (Buschmann et al., 2015). However, AIR9 is substantially different from MASP1 in that its microtubule binding region is at the N-terminal portion of the protein rather than the C-terminus as in MASP1 and the remainder of AIR9 outside of the N-terminal LRR-repeat region is divergent from, and much larger than, MASP1. We have not observed MASP1 localization on spindle fibers or newly forming cell plate and the cellular function of MASP1 remains unclear.

Interestingly, it has been reported that *EGR*s have a gradient of expression across the root meristem. Sorting of root meristem cells into proximal (close to QC), medial, and distal regions based on GFP expression driven by the *Plant U-Box25* (*PUB25*) or *SPATULA* (*SPT*) promoters found that *EGR1, EGR2* and *EGR3* had low expression in the proximal meristem region but significantly higher expression in medial and distal meristem regions (Wendrich et al., 2017). While this is an intriguing observation, it is not known whether such gradient in EGR gene expression also leads to differences in EGR protein level in the proximal versus distal meristem regions and whether spatial differences in EGR protein level are functionally important for growth regulation.

Here we demonstrate that differences in EGR-MASP1 stoichiometry in different regions of the root meristem controls root meristem size and activity during low ψ_w_. These results show that EGRs and MASP1 link stress signaling with regulation of root meristem function during drought stress and show how disruption of EGR-MASP1 stoichiometry and signaling can enhance growth maintenance, or further downregulate growth, during moderate severity low ψ_w_. We also found that EGR2 was most highly expressed in root cortex cells, an observation that is interesting in light of recent data indicating a key role of the cortex in regulating growth responses to low ψ_w_.

## Results

### EGRs and MASP1 have opposing effects on root elongation during low ψ_w_

To further determine how EGRs and MASP1 affect root growth responses to low ψ_w_, we transferred 5-day-old seedlings to PEG-agar plates of moderate severity low ψ_w_ stress (−0.7 MPa). This ψ_w_ was selected as we have previously shown that it reduces growth by a moderate amount (50-70 percent) in both PEG-agar plate assays and soil drying experiments (Bhaskara et al., 2017; Wong et al., 2019) and is within the range of soil ψ_w_ that occurs during moderate severity drought in many types of field environments. In these experiments, it was visually clear that *egr1-1egr2-1* and *35S:MASP1* maintained higher root and shoot growth compared to wild type at low ψ_w_ (Fig. 1A), consistent with previous results where whole seedling fresh and dry weights were quantified (Bhaskara et al., 2017). Quantitation of root elongation rates found that in wild type root elongation rate was reduced by nearly 70 percent compared to the unstressed control over six days after transfer to low ψ_w_ (Fig. 1B). The *egr1-1egr2-1* double mutant maintained significantly higher root elongation rates than wild type at low ψ_w_ over this period (Fig. 1B). We also checked root elongation rates of *egr* single mutants and found that there was a trend of increased root elongation at low ψ_w_ for *egr1* and *egr2* mutants (but the difference was only significant for *egr2-1*) while *egr3* mutants had similar or lesser effect (Supplemental Fig. S1A). Together, these data confirmed that *EGR* phosphatases act redundantly to restrict root elongation during low ψ_w_ stress.

**Figure 1:**
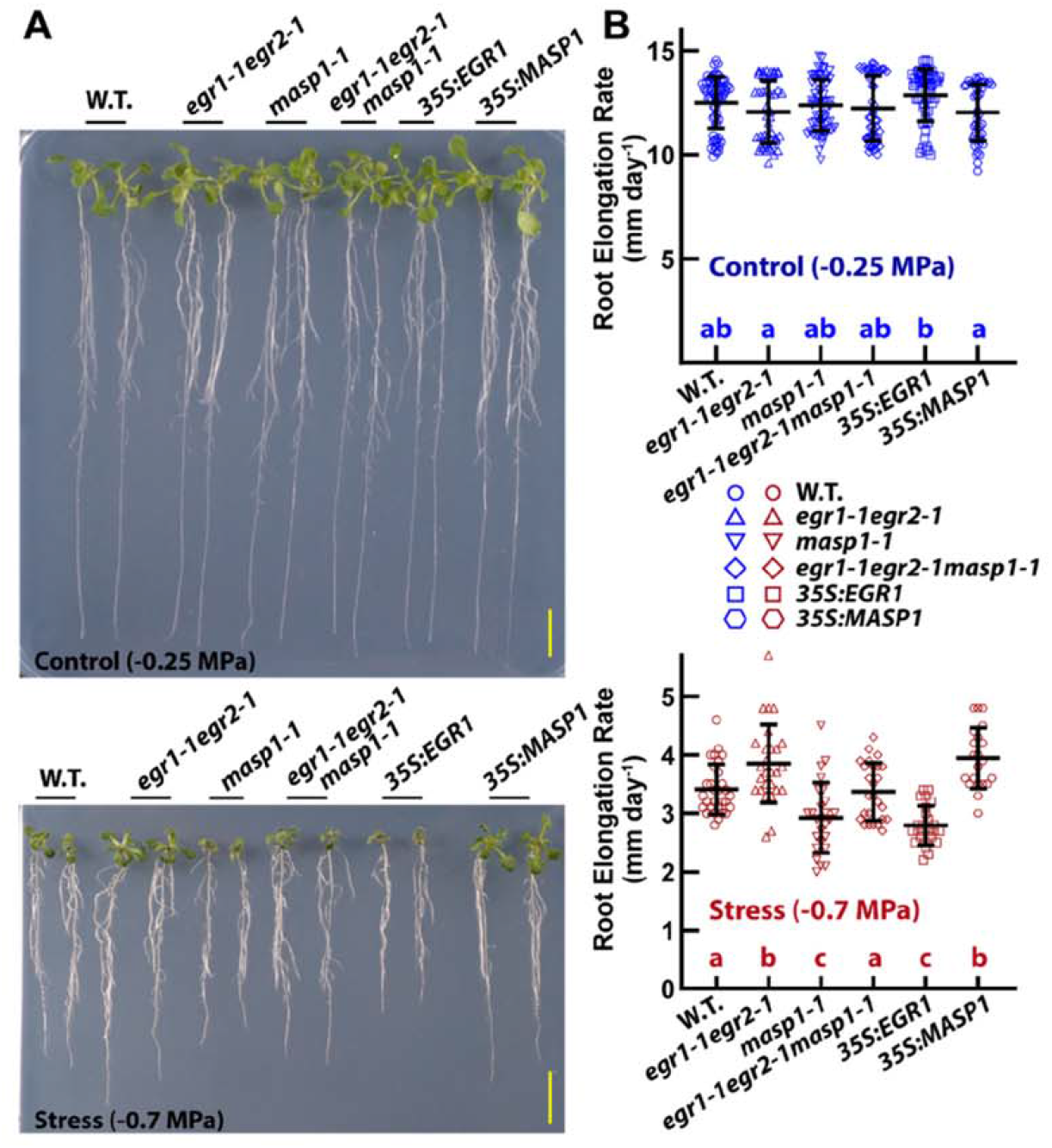
EGRs and MASP1 affect growth during low ψ_w_ stress. A. Representative seedlings of *EGR* and *MASP1* mutants and overexpression lines. Five-day-old seedlings were transferred to either fresh control plates (−0.25 MPa) or PEG-infused agar plates for moderate severity low ψ_w_ (−0.7 MPa) treatment. Plants were photographed at the end of the experiment (six days after transfer). Scale bars indicate 1 cm. B. Primary root elongation rates over six days after transfer of seedlings to control or stress (−0.7 MPa) treatments. Blue or red symbols show elongation rates for individual seedlings. Black bars and error bars indicate the mean and standard deviation for each genotype (n = 22 to 67). Lower case letters in the stress panel indicate statistical differences (ANOVA with Tukey’s post hoc test, corrected P ≤ 0.05). Genotypes sharing the same letter did not significantly differ from one another. Data are combined from three independent experiments and three independent transgenic lines per construct. Both *35S:EGR1* and *35S:YFP-EGR1* as well as *35S:MASP1* and *35S:YFP-MASP1* lines were used. Data from additional experiments with each individual transgenic line is shown in Supplemental Fig. S2. The lines expressing untagged or YFP-tagged proteins had consistent result and the combined data are labeled as *35S:MASP1* and *35S:EGR1* for clarity of presentation.

Ectopic *MASP1* expression (*35S:MASP1*) led to higher root elongation rate at low ψ_w_ compared to wild type while *masp1-1* and *35S:EGR1* had reduced root elongation rate (Fig. 1B). This also was consistent with previous quantitation of seedling dry weights in *masp1-1* (and *masp1-2*) as well as *35S:EGR1* which found that all these genotypes had decreased growth at low ψ_w_ but had little effect on growth in the unstressed control (Bhaskara et al., 2017). For the analyses presented in Fig. 1 we used transgenic lines with expression of untagged EGR1 or MASP1 and lines with expression of either protein with an N-terminal fusion to YFP. Transgenic lines used in this study are listed in Supplemental Table S1 and additional data showing that individual transgenic lines have similar effect on root elongation is shown in Supplemental Fig. S2. Note that *35S:EGR1* and *35S:MASP1*, as well as constructs with an N-terminal YFP added to either protein, have been previously shown to complement their respective mutants (Bhaskara et al., 2017). Lines expressing either tagged or untagged EGR1 and MASP1 were used in this study to ensure that our results were robust across multiple independent transgenic lines and as an extra check that the presence of the YFP-tag did not influence the results so that later experiments examining the expression patterns of YFP-tagged EGR1 and MASP1 could be robustly compared to the results of physiology experiments. As all EGR1 and MASP1 transgenic lines used in this study had consistent phenotypes, for subsequent experiments combined data of multiple transgenic lines is presented in the main text figures and the lines referred to as *35S:EGR1* and *35S:MASP1* for clarity in presentation while data for individual transgenic lines are presented in Supplemental Figures.

Interestingly, the *egr1-1egr2-1masp1-1* triple mutant was intermediate between *egr1-1egr2-1* and *masp1-1* and did not significantly differ from wild type at low ψ_w_ (Fig. 1B). This intermediate phenotype of *egr1-1egr2-1masp1-1* was consistent with our previous observations and our interpretation that MASP1 acts downstream of EGRs; but, EGRs also have MASP1-independent effect(s) on growth (Bhaskara et al., 2017).

We also measured growth of EGR-MASP1 mutants and transgenic lines after transfer to media containing NaCl. Salt stress has both an osmotic component and a sodium toxicity component (Munns and Tester, 2008). To avoid excessive sodium toxicity which may obscure the osmotic stress response, we used relatively mild salt stress treatments (75 mM and 125 mM) which reduced root elongation rate of wild type by 30 percent and 50 percent, respectively (Supplemental Fig. S3A). These experiments showed that the EGR-MASP1 mutant and transgenic lines exhibited similar differences in root elongation during salt stress as during low ψ_w_, albeit that the effect sizes were smaller because root elongation rates of all genotypes remained relatively high. At 75 mM NaCl, *egr1-1egr2-1* and *35S:MASP1* had significantly higher root elongation rates than the contrasting *masp1-1* and *35S:EGR1* genotypes, respectively. At 125 mM NaCl, *egr1-1egr2-1* and *35S:MASP1* had higher root elongation rates than wild type or other genotypes. Along with previous data showing that EGRs and MASP1 affect rosette growth during soil drying (Bhaskara et al., 2017), these data demonstrate that the effects of EGR and MASP1 on growth can be observed in multiple treatments that restrict water availability.

### MASP1 promotes cell division during low ψ_w_ and is required for the increased cell division of *egr1-1egr2-1*

The effect of EGR and MASP1 on root growth at low ψ_w_ could be caused by a change in the number of cells produced by cell division, change in cell size or a combination of the two. To analyze cell division, we first used a reporter line containing β-glucuronidase (GUS) fused to the Cyclin B1 promoter and cyclin destruction box (CYCB1::GUS) (Colon-Carmona et al., 1999). This reporter produces foci of GUS activity when a cell goes through the G2-M phase transition thus allowing mitotic activity to be estimated by counting the number GUS foci. In wild type, low ψ_w_ reduced the number of CYCB1::GUS foci by more than half; consistent with many previous reports that low ψ_w_ decreases cell division (Fig. 2A and B; data from individual transgenic lines is shown in Supplemental Fig. S4A). In the low ψ_w_ treatment, *egr1-1egr2-1* had increased number of GUS foci compared to wild type. *35S:MASP1* had increased numbers of CYCB1::GUS foci in both the low ψ_w_ and unstressed control treatments; however, the effect was proportionally much larger in the low ψ_w_ treatment. Interestingly, the *egr1-1egr2-1masp1-1* triple mutant had the same low level CYCB1::GUS as *masp1-1* (Fig. 2A and B). Thus, the increased cell division of *egr1-1egr2-1* at low ψ_w_ was dependent upon MASP1. Note that the CYCB1::GUS reporter does not allow one to determine which cell layer the GUS foci are located in. This does not affect interpretation of our results as none of our mutants or transgenic lines disrupted the arrangement of different cell types within the root tip (see below).

**Figure 2:**
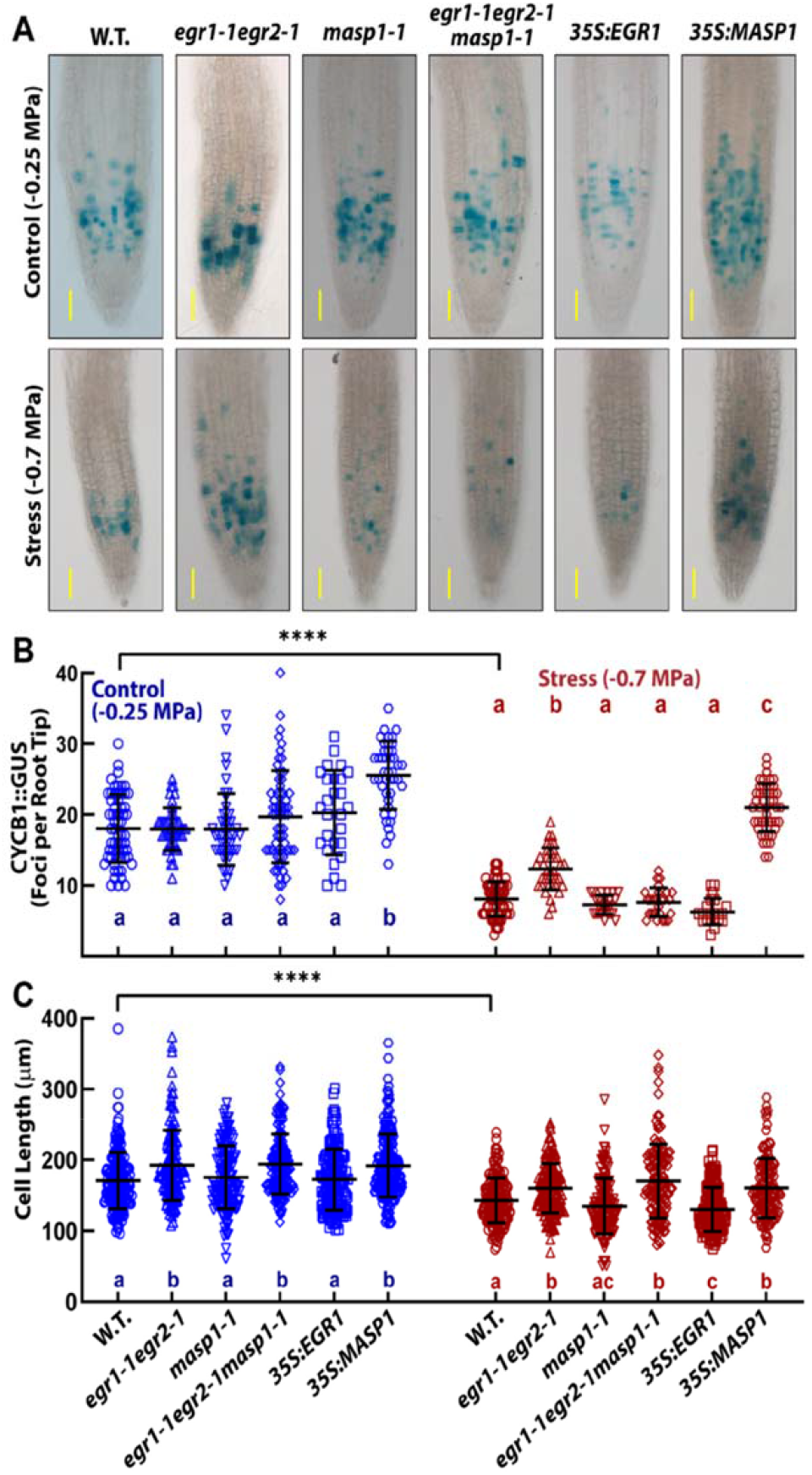
EGRs and MASP1 affect root cell division and cell expansion during low ψ_w_ stress. A. Representative images of CYCB1::GUS staining in primary root tips of the indicated mutant and ectopic expression genotypes 4 days after transfer of five-day-old seedlings to either control or low ψ_w_ (−0.7 MPa). Scale bars indicate 100 μm. B. Quantification of CYCB1::GUS foci in the primary root meristem. Symbols indicate counts of GUS foci from individual roots while black bars and error bars indicate the mean and standard deviation for each genotype. Within the control or stress treatments, genotypes sharing the same letter do not significantly differ from one another (ANOVA with Tukey’s post hoc test, corrected P ≤ 0.05). In wild type, the number of CYCB1::GUS foci was significantly different between control and stress treatment (T-test, P ≤ 0.001). Data are combined from three or four independent experiments (n = 21-66) for each genotype and at least two independent transgenic lines for *35S:MASP1* and *35S:EGR1*. Data plotted separately for each transgenic line is shown in Supplemental Fig. S4. C. Cell length at 2 mm from the root tip measured four days after transfer of mutant or ectopic expression seedlings to control or low ψ_w_ (−0.7 MPa). Format of data and statistical analysis are the same as in B (n = 133-224). Both *35S:EGR1* and *35S:YFP-EGR1* as well as *35S:MASP1* and *35S:YFP-MASP1* lines were used. The lines expressing untagged or YFP-tagged proteins all had consistent result and the combined data are labeled as *35S:MASP1* and *35S:EGR1* for clarity of presentation. Data plotted separately for each transgenic line is shown in Supplemental Fig. S4.

The CYCB1::GUS reporter can also be induced by DNA damage (Schnittger and De Veylder, 2018). This is unlikely to have affected our results as the moderate severity stress used here is not expected to cause DNA damage and we observed reduced CYCB1::GUS under low ψ_w_ rather than increased activity that would be expected if substantial DNA damage occurred. Nonetheless, we also assessed cell division activity using 5-ethynyl-2′-deoxyuridine (EdU) staining to label newly synthesized DNA. Compared to the CYCB1::GUS results, higher numbers of EdU foci were observed in the meristem region (apical 300 μm of the root) for both control and stress treatments (Supplemental Fig. S5). This is likely because of the amount of time needed to effectively stain the roots and because the EdU stain is not actively removed after DNA replication is complete. Despite this difference, it was clear that the number of EdU foci was reduced by approximately half in low ψ_w_-treated wild type compared to the unstressed control. Also consistent with the CYCB1::GUS results, *egr1-1egr2-1* and *35S:MASP1* had significantly increased EdU staining compared to wild type at low ψ_w_ while *masp1-1, egr1-1egr2-1masp1-1* and *35S:EGR1* were low but not significantly different from wild type (Supplemental Fig. S5). This further indicated that EGRs suppressed cell division at low ψ_w_ while MASP1 promoted it. In EdU staining of the unstressed control, none of the mutant or ectopic expression lines differed from wild type (Supplemental Fig. S5).

### EGRs affect cell size independently of MASP1

In the Arabidopsis primary root, cells are fully expanded once they reach 1.5 to 2 mm from the root apex (van der Weele et al., 2000; West et al., 2004; Yang et al., 2017). Therefore, we measured epidermal cell length at 2 mm from the root apex (Fig. 2C). Cell length of unstressed wild type was similar to previous reports (West et al., 2004; Yang et al., 2017). Also consistent with previous reports, we found that root cell expansion was less affected by low ψ_w_ than cell division. In wild type, low ψ_w_ reduced cell length by 20 percent compared to the unstressed control. Interestingly, *egr1-1egr2-1* and *egr1-1egr2-1masp1-1* had increased cell size in both the stress and control treatments while *masp1-1* did not. Thus, loss of EGR activity led to enhanced cell expansion and, in contrast to cell division, the effect of *egr1-1egr2-1* on cell expansion did not depend on MASP1. These data explain the intermediate growth phenotype of *egr1-1egr2-1masp1-1* at low ψ_w_ as the triple mutant had the increased cell expansion of *egr1-1egr2-1* but not the increased cell division. We also found that the *egr* single mutants tended to have slightly larger cells than wild type under low ψ_w_; however, there was not a significant difference for any of the individual mutants (Supplemental Fig. S1B). This indicated that *EGR1* and *EGR2* act redundantly to restrict cell expansion during low ψ_w_ stress.

Conversely, *35S:EGR1* further decreased cell size by a small amount under low ψ_w_ but had no effect in the unstressed control, consistent with the overall strong growth restriction of *35S:EGR1* compared to wild type at low ψ_w_ and lack of *35S:EGR1* effect on unstressed plants (Fig. 1A; data for individual transgenic lines is shown in Supplemental Fig. S4B). Despite its decreased growth at low ψ_w_, *masp1-1* did not have significantly smaller cells; although, this was due in part to variability of cell size in *masp1-1*, as we also noted in previous examination of hypocotyl cells (Bhaskara et al., 2017). *35S:MASP1* had slightly bigger cells in both control and stress treatments, similar to *egr1-1egr2-1*. As the regulation of cell size and cell division involves both coordination and compensatory mechanisms (Gonzalez et al., 2012; Clauw et al., 2016), some of the differences in cell size observed may be indirect effects of changes in cell division or meristem size.

### *egr1-1egr2-1* and *35S:MASP1* maintain a larger root meristem at low ψ_w_

To determine whether the effects of EGRs and MASP1 on cell division may alter cellular organization within the root meristem, we examined the proximal root meristem region just behind the QC for all genotypes in control and low ψ_w_ stress treatments (Fig. 3A). The QC region as well as epidermal, cortex and endodermal cell layers were intact in all genotypes and there was no evidence of improperly placed cell divisions. In the stress treatment, epidermal, cortex and endodermal cells in the proximal meristem region were larger (longer) than they were in unstressed roots (Fig. 3A). This was consistent with data in Figure 2 showing that low ψ_w_ inhibited cell division to greater extent than cell expansion. Cell shape also became more irregular at low ψ_w_, perhaps associated with the altered coordination between cell division and cell expansion. The larger meristem cell size was particularly apparent in *egr1-1egr2-1masp1-1* but was less pronounced in *egr1-1egr2-1* and *35S:MASP1*, consistent with the different effects of these genotypes on cell division in the CYCB1::GUS assays.

**Figure 3:**
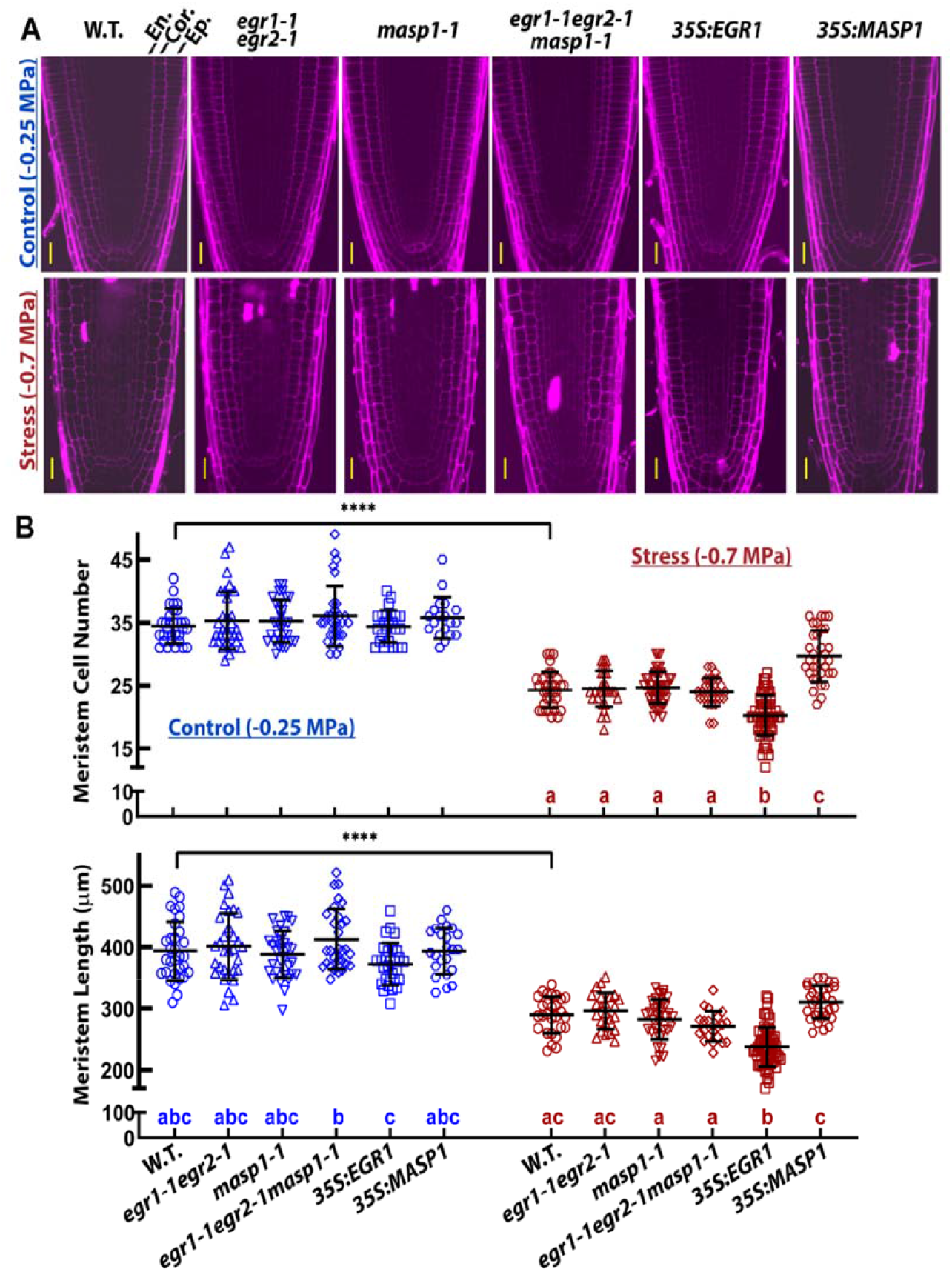
Ectopic EGR or MASP1 expression alters root meristem size during low ψ_w_ stress. A. Representative images of PI stained root tips to show the cell size and organization of cell files in the proximal meristem region. Scale bars indicate 20 μm. The epidermal (Ep.), cortical (Cor.) and endodermal (En.) cell layers are labeled on the wild type control image. B. Meristem cell number and meristem length five days after transfer of five-day-old mutant and ectopic expression seedlings to unstressed control or to low ψ_w_ (−0.7 MPa). Data are combined from 2-3 independent experiments for each genotype (n = 19-31 in the control, n = 33-65 in the stress treatment). Symbols indicate data from individual roots while black bars and error bars indicate the mean and standard deviation for each genotype. Within the stress treatments, genotypes sharing the same letter do not significantly differ from one another (ANOVA with Tukey’s post hoc test, corrected P ≤ 0.05). There were no significant differences among genotypes for meristem cell number in the unstressed control. In wild type, meristem size was significantly different between control and stress treatment (T-test, P ≤ 0.001). In the stress treatment, the increase of meristem length in *35S:MASP1* was marginally non-significant compared to wild type (adjusted P = 0.08) and *egr1-1egr2-1* may have a slightly longer meristem than *egr1-1egr2-1masp1-1* (adjusted P = 0.06). Representative images showing differences in root meristem length are shown in Supplemental Fig. S6). For *35S:EGR1* and *35S:MASP1*, three independent transgenic lines were used for each gene. Data for each individual transgenic line plotted separately is shown in Supplemental Fig. S7.

To quantify EGR and MASP1 effects on root meristem size, we counted the number of cells in a file of epidermal cells from the QC to the first elongated cell of the transition zone and also measured the distance from the QC to the first elongated cell (Fig. 3B, representative images to illustrate differences in meristem length are shown in Supplemental Fig. S6 and data for individual transgenic lines is shown in Supplemental Fig. S7). Meristem cell number and size was significantly reduced by low ψ_w_ in wild type, similar to previous observations in seedlings exposed to moderate severity salt stress (West et al., 2004). Compared to wild type, the number of cells in the root meristem was increased in *35S:MASP1* but decreased in *35S:EGR1* at low ψ_w_ (Fig. 3A; Supplemental Fig. S7), consistent with their effects on growth and cell division. Meristem length was also strongly decreased in *35S:EGR1* at low ψ_w_. Conversely, meristem length of *35S:MASP1* at low ψ_w_ was significantly larger than that of *masp1-1* or *egr1-1egr2-1masp1-1*. The *egr* and *masp1* mutants did not significantly differ from each other or from wild type in meristem cell number or meristem length at low ψ_w_ (although *egr1-1egr2-1* may have a slightly longer meristem than *egr1-1egr2-1masp1-1*, adjusted P = 0.06). Also, there were no significant differences in meristem size among genotypes in the unstressed control. Similar effects on root meristem size were seen at four days after transfer to low ψ_w_ (Supplemental Fig. S8). In seedlings transferred to 75 mM NaCl, *egr1-1egr2-1* and *35S:MASP1* had higher meristem cell number and larger meristem size than *masp1-1* or *35S:EGR1* (Supplemental Fig. S3B).

Roots of all genotypes were thinner after 5 days at low ψ_w_ compared to unstressed roots (Supplemental Fig S6). This was consistent with previous observations that low ψ_w_ prevented the increase in root diameter that occurs over time in unstressed Arabidopsis roots (van der Weele et al., 2000). *35S:MASP1*, and perhaps *egr1-1egr2-1,* maintained a higher root diameter, while *masp1-1* was thinner. This may be consistent with the interpretation of van der Weele et al. (2000) that Arabidopsis root diameter was positively correlated with root elongation rate. Root thinning in response to low ψ_w_ has also been observed in a number of other plant species, such as maize (Sharp et al., 1988).

As the root meristem is small in newly germinated seeds and later increases in size as root growth accelerates (Beemster and Baskin, 1998; Biancucci et al., 2015) we also examined root meristem size in younger seedlings. In 2-and 4-day-old unstressed seedlings, *egr1-1egr2-1masp1-1* had increased meristem size compared to other genotypes (Supplemental Fig. S9). This indicated that EGRs and MASP1 may have complementary roles in controlling root meristem size during early seedling development in addition to their roles in regulating meristem size under low ψ_w_ stress.

### Spatial differences in EGR2 versus MASP1 stoichiometry influence meristem size during low ψ_w_ stress

Examination of EGR2 and MASP1 root tip expression pattern illustrated how they can interact to regulate meristem size and also indicated why ectopic EGR or MASP1 expression had such dramatic effects on root meristem cell number and meristem size. Among the three EGR PP2Cs, *EGR2* was selected for *promoter:GUS* analysis because EGR2 has relatively high expression and because *egr2* single mutants had slightly larger effect on growth compared to other *egr* single mutants (Supplemental Fig. S1; Bhaskara et al., 2017; Wendrich et al., 2017). In unstressed plants, *EGR2_promoter_:GUS* expression was high in the mature region but undetectable in the apical 1 mm of the root tip which included the meristem and cell expansion regions (Fig. 4A). Low ψ_w_ stress led to an increase in *EGR2_promoter_:GUS* expression including detectable expression closer to the root tip. This may be related to a shifting of the cell expansion and cell differentiation zones closer to the root tip, as has been observed for high temperature and other treatments that inhibit root elongation (Yang et al., 2017). However, *EGR2_promoter_:GUS* activity was still not detected in the apical 200-300 μm which contains the root meristem. *MASP1_promoter_:GUS* expression was essentially the converse of *EGR2* with high expression in the proximal meristem region just behind the QC in both control and stress treatments (Fig. 4A). Low ψ_w_ induced a higher level of *MASP1_promoter_:GUS* staining across the root meristem as well as an enhanced, but still relatively low, level of expression in mature root tissue. This was consistent with our previous observation that *MASP1* gene expression and protein level of whole seedlings increased in plants exposed to a more severe −1.2 MPa low ψ_w_ treatment (Bhaskara et al., 2017).

**Figure 4:**
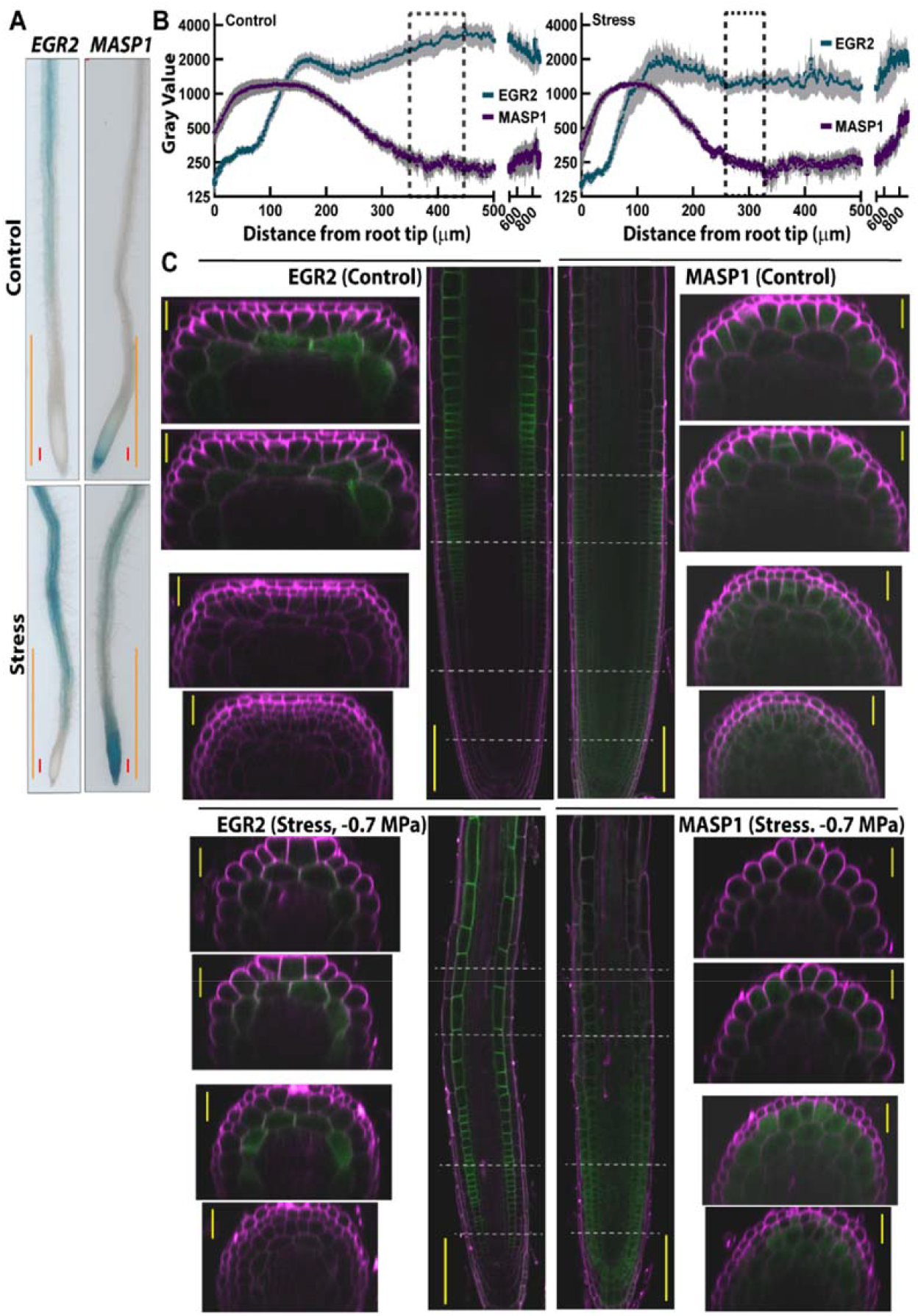
EGR and MASP1 have differing spatial patterns of expression in the root tip. A. Representative images of *EGR2_promoter_:GUS* and *MASP1_promoter_:GUS* staining in primary root tips four days after transfer to control or low ψ_w_ (−0.7 MPa). Red scale bars indicate 100 μm. Orange bars indicate 1 mm. Images are representative of the expression patterns seen in three independent transgenic lines for each construct. B. Quantitation of *EGR2_promoter_:EGR2-YFP* and *MASP1_promoter_:MASP1-YFP* fluorescence intensity along the root tip. Roots were imaged with a fluorescence imager and Image J was used to perform line scans of root tips for 6-10 roots (from two independent transgenic lines for each construct assayed in two independent experiments) at four days after transfer of seedlings to control or low ψ_w_ (−0.7 MPa) treatments. Representative root tip images used for the scans are shown in Supplemental Fig. S12C. Dashed line boxes indicate the end of the root meristem for control and stress treatments (based on data in Fig. 3B). Gray error bars (gray shading) indicate the standard error. Note the log scale of the y-axis. C. Root tip expression pattern of *EGR2_promoter_:EGR2-YFP* and *MASP1_promoter_:MASP1-YFP* in unstressed plants (Control) and low ψ_w_-treated plants (−0.7 MPa, 4 days). The gray dashed lines in the whole root tip images indicate the positions of the radial scans (50, 150, 350 and 450 μm from the quiescent center). The whole root tip images were made by merging individual images using Leica LAS X software. Images are representative of the expression pattern seen in three independent transgenic lines for each construct. In the whole root tip images, scale bars indicate 100 μm. In the radial scan images, scale bars indicate 20 μm.

To examine patterns of EGR2 and MASP1 protein accumulation, we generated plants transformed with a genomic fragment containing the *EGR2* or *MASP1* promoter and gene body with C-terminal fusion to YFP. We demonstrated that both *MASP1	:MASP1-YFP* and *EGR2_promoter_:EGR2-YFP* could complement the growth phenotypes of *masp1-1* or *egr1-1egr2-1* at low ψ_w_ (Supplemental Fig. S10A and S11A). Note that *egr1-1egr2-1* transformed with *EGR2_promoter_:EGR2-YFP* still had a small increase in growth compared to wild type, again consistent with redundant function of EGR1 and EGR2 in growth regulation at low ψ_w_ (Supplemental Fig. S11A).

A broad scale analysis using line scans of fluorescence intensity along the root tip for *EGR2_promoter_:EGR2-YFP* and *MASP1_promoter_:MASP1-YFP* constructs showed that EGR2 was very low in the apical 50-100 μm of the root but increased nearly 10-fold in more distal parts of the root tip (Fig. 4B). In contrast MASP1 protein level was highest in the apical 100 μm but declined thereafter to a low basal level. MASP1 protein was detected at positions farther from the root tip than *MASP1_promoter_:GUS* expression, which was highly concentrated in the proximal root meristem of unstressed plants. This was consistent with cells being displaced away from the root tip as the MASP1 protein was being synthesized. In low ψ_w_-treated roots both the decrease in MASP1 and increase in EGR2 occurred closer to the root tip (Supplemental Fig. S12A, Fig. 4B). The point at which MASP1 expression reached a minimum value corresponded to the end of the root meristem under both control and low ψ_w_ conditions (indicated by the dashed boxes in Fig. 4B). Interestingly, MASP1 protein level did not substantially increase in the root meristem under low ψ_w_ stress despite the strong increase in *MASP1_promoter_:GUS* activity at low ψ_w_ (compare Fig. 4A and B). The higher *MASP1_promoter_:GUS* activity at low ψ_w_, along with reduced root elongation and slower displacement of cells away from the root tip would be expected to lead to higher MASP1 protein level. Since this was not observed, it is possible that MASP1 protein stability is affected by low ψ_w_ to control its abundance and spatially restrict MASP1 protein accumulation. Conversely EGR2 accumulated to relatively high level in the region 200-400 μm from the root tip despite minimal EGR2 promoter activity in this region (compare Fig. 4A and B), suggesting that EGR2 is a relatively stable (or efficiently translated) protein compared to MASP1.

We also conducted a similar line scan analysis of root tips from plants expressing *35S:YFP-EGR1* and *35S:YFP-MASP1* (Supplemental Fig. S12B). The *35S* promoter led to a relatively even protein level across the root tip for both EGR1 and MASP1. This meant that *35S*-driven EGR1 expression was more than 10-fold higher than endogenous EGR2 in the apical 50-80 μm of the root. Conversely, *35S*-driven MASP1 expression was higher than endogenous MASP1 in the region 200-500 μm from the root tip. These data indicated that *35S:EGR1* and *35S:MASP1* strongly affect root meristem size and growth at low ψ_w_ because *35S*-driven ectopic expression of either gene disrupts spatial patterns of EGR versus MASP1 stoichiometry in the root tip. This was supported by observation that, in contrast to *35S:EGR1*, expression of *EGR2_promoter_:EGR2-YFP* in the wild type background did not inhibit growth at low ψ_w_ (Supplemental Fig. S11A) and observation that *MASP1_promoter_:MASP1-YFP* in the wild type background did not promote growth at low ψ_w_ (Supplemental Fig. S10A). It was also interesting to note that root tip expression of *35S*-driven EGR1 was similar in both control and stress conditions (Supplemental Fig. S12B) despite the fact that *35S:EGR1* plants only had reduced root growth, cell division or cell size in the stress treatments (Fig, 1, 2, 3; Supplemental Fig. S3). This raises the possibility that a stress-related factor is required to potentiate the effect of ectopic EGR expression.

High resolution imaging of the root tip showed that EGR2 protein level was highest in the cortical cell layer with much lower level of EGR2 expression in some epidermal cells (Fig. 4C, Supplemental Fig. S11C). In unstressed roots, EGR2 was absent from the proximal meristem region but there was a peak of EGR2 in the distal meristem region. Observations in multiple transgenic lines found that this peak of EGR2 always preceded the end of the cell division zone, even though its exact position could vary from root to root. A similar pattern was observed in roots exposed to low ψ_w_; however, we consistently observed that low ψ_w_ caused EGR2 expression to encroach closer to the QC (Fig. 4C; Supplemental Fig. S11C). Radial scans at in the proximal meristem region (50 and 150 μm from the QC) and distal meristem to beginning of the cell expansion zone (350 and 450 μm from the QC) confirmed the gradient in EGR2 protein level and also confirmed that EGR2 was mainly expressed in cortical cells (Fig. 4C; positions of the radial scans are indicated by dashed lines on the root tip images). Note however that EGR2 was sometimes difficult to fully visualize in radial scans unless the scan was sufficiently close to a transverse side of a cortex cell where EGR2 signal was highest.

Conversely, MASP1 protein level of unstressed roots was high in the region just behind the QC and was expressed across all cell types. In the distal meristem and transition to cell elongation (approximately 250 to 450 μm behind the QC), MASP1 protein level declined in the inner cell layers (including cortex) but could still be detected in epidermal cells into the cell elongation zone (Fig. 4C; Supplemental Fig. S10B). At low ψ_w_, we consistently observed that MASP1 protein level remained high in the proximal meristem region but declined substantially in the distal meristem and beginning of the cell expansion region (Fig. 4C; Supplemental Fig. S10B). Radial scans confirmed this decline of MASP1 protein level and showed that it occurred across all cell types (Fig. 4C). These data demonstrated that in unstressed plants EGR2 and MASP1 overlapped in the root cortex near the distal end of the root meristem. Low ψ_w_ stress increased the EGR2-MASP1 overlapping region and pushed it closer to the QC, consistent with shortening of the root meristem in response to low ψ_w_.

### Phosphomimic MASP1 (S670D), but not wild type MASP1, can overcome *35S:EGR1* suppression of root meristem size at low ψ_w_

Previously we demonstrated that EGRs (either directly or indirectly) attenuate MASP1 phosphorylation and that phosphonull MASP1^S670A^ was inactive and unable to complement the *masp1-1* mutant (Bhaskara et al., 2017). Consistent with these previously reported growth phenotypes, we found that *35S:MASP1^S670D^* increased meristem size while *35S:MASP1^S670A^* slightly decreased meristem size under low ψ_w_ (Fig. 5A and B). Combined with the results presented above, this suggested that EGR control of MASP1 S670 phosphorylation in their overlapping zone of expression is a key factor controlling root meristem size during low ψ_w_. If this hypothesis is correct, we expect that *35S:EGR1* will be able to suppress the increased meristem size of *35S:MASP1* plants but would be unable to suppress meristem size in plants expressing *35S:MASP1^S670D^*. To test this hypothesis, we crossed both wild type and *35S:EGR1* to *35S:MASP1*, *35S:MASP1^S670D^* and *35S:MASP1^S670A^* and then examined root meristem size of low ψ_w_-treated F_1_ seedlings. Consistent with our hypothesis, F_1_ seedlings of *35S:MASP1* x W.T. had significantly larger root meristem size than W.T. and larger than F_1_ seedlings of *35S:MASP1* x *35S:EGR1* (Fig. 5C). In contrast, F1 seedlings of *35S:MASP1^S670D^* x *35S:EGR1* had the same increase in meristem size as *35S:MASP1^S670D^* x W.T., indicating that *35S:EGR1* was unable to counteract the effect of phosphomimic MASP1. *35S:MASP1^S670A^* was again ineffective and did not increase meristem size in F_1_ seedlings of the W.T. or *35S:EGR1* crosses (Fig. 5C). These data supported our hypothesis that EGR control of MASP1 S670 phosphorylation is a key mechanism to control root meristem size during low ψ_w_ stress.

**Figure 5:**
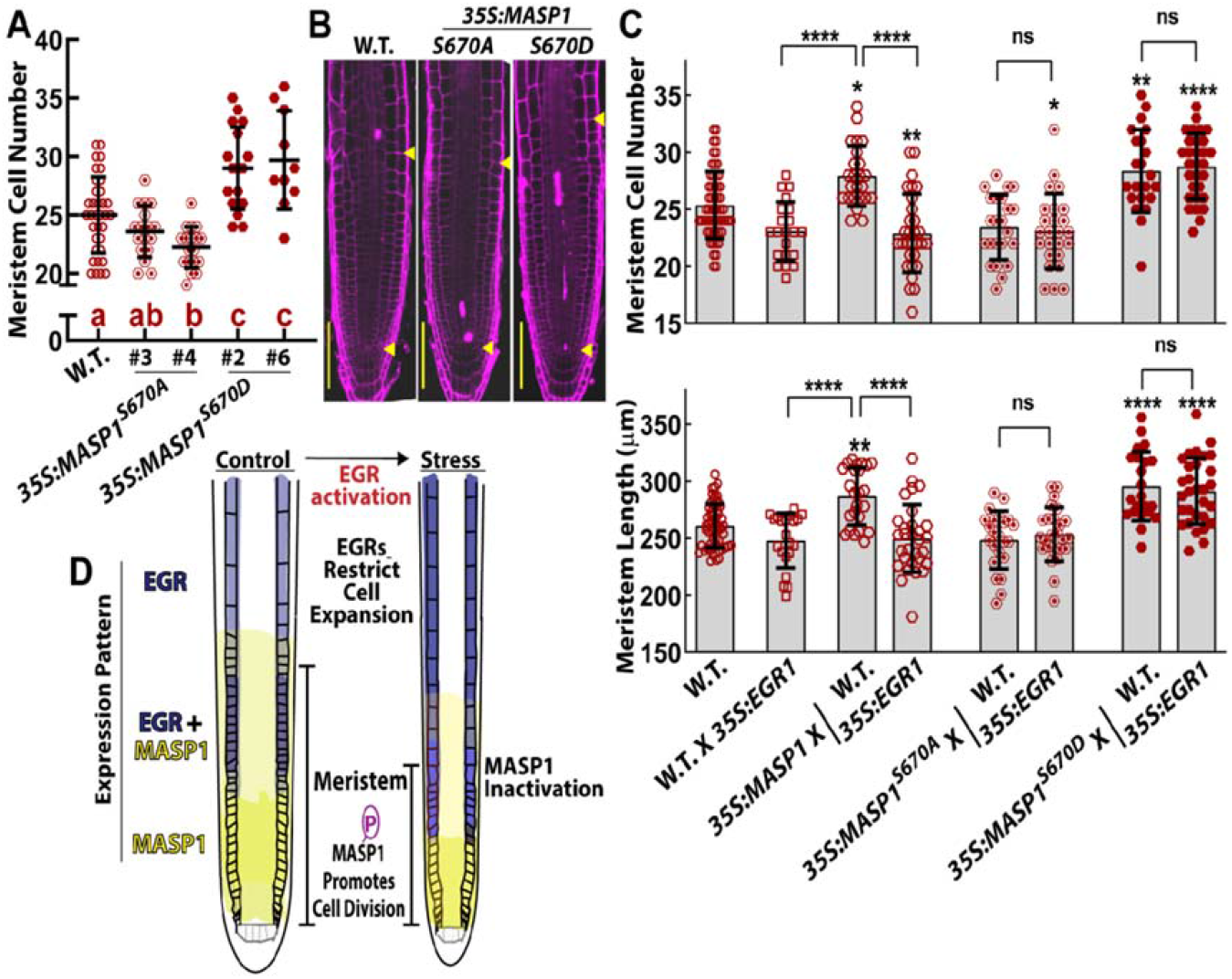
EGR attenuation of MASP1 S670 phosphorylation in their zone of overlapping expression controls root meristem size at low ψ_w_. A. Root meristem cell number at low ψ_w_ for wild type and two independent transgenic lines expressing phosphonull MASP1 (*35S:MASP1^S670A^*) or phosphomimic MASP1 (*35S:MASP1^S670D^*). The MASP1 phosphonull and phosphomimic lines were previously described in Bhaskara et al. (2017). Data are combined from 2-3 independent experiments (n = 35 for wild type and 10-18 for each transgenic line). The transgenic lines used here are in the GFP-TUA6 background (Supplemental Table S1). It was previously shown that presence of GFP-TUA6 did not affect the growth phenotypes of any EGR-MASP1 mutant or transgenic lines (Bhaskara et al., 2017). Symbols indicate data from individual roots while black bars and error bars indicate the mean and standard deviation for each genotype. Different letters indicate significant differences between genotypes (ANOVA with Tukey’s post hoc test, corrected P ≤ 0.05). B. Representative root meristem images of each genotype in A. Scale bars indicate 100 μm. C. Quantitation of meristem size in F_1_ seedlings of *35S:EGR1* crossed to lines ectopically expressing non-mutated MASP1, phosphonull MASP1 (S670A) or phosphomimic MASP1 (S670D). All the MASP1 lines express untagged MASP1 in the GFP-TUA6 background. Root meristem size was measured five days after transfer to low ψ_w_ (−0.7 MPa). Bars indicate the mean, error bars indicate the standard deviation from 2 independent experiments for each genotype (n = 56 for wild type, n = 21-29 for F_1_ genotypes). Asterisks directly above the bars indicate a significant difference compared to wild type, brackets show the results of comparison between the indicated genotypes (ANOVA with Tukey’s post hoc test, * indicates corrected P ≤ 0.05; ** P ≤ 0.005; *** P ≤ 0.0001). Note that the difference in meristem cell number between wild type (W.T.) and W.T. x *35S.EGR1* was marginally nonsignificant (adjusted P = 0.07). D. Model of how EGR-MASP1 signaling affects meristem function and root growth at low ψ_w_. MASP1 is highly-expressed in all cell types of the proximal root meristem where it promotes cell division. MASP1 decreases in expression farther from the QC. especially in the inner cell layers and cortex. Conversely, EGR2 is low in the proximal meristem but high in the distal meristem and elongation zone where it influences cell expansion. EGR2 is highly expressed in cortical cells while MASP1 docs not have cell type specific expression. During low ψ_w_ stress. EGR2 expression encroaches closer to the proximal meristem region. In their overlapping zone of expression. EGRs attenuate phosphory lation of MASP1, particularly in cortex cells, to suppress MASP1 activity and restrict meristem size. The high expression of EGR2 in cortex is consistent with recent data indicating that this cell layer is important for growth responses to low ψ_w_. Low ψ_w_ activates (or de-represses) EGRs such that EGR-MASP1 signaling is a dominating factor controlling root elongation and meristem function during low ψ_w_ stress but has less or no effect on unstressed plants.

### EGR-MASP1 shoot expression patterns

EGRs and MASP1 also clearly affect shoot growth (Fig. 1A; Bhaskara et al., 2017) and EGR2-MASP1 expression patterns suggested that a similar mechanism of opposing EGR-MASP1 signaling may also operate in shoot and leaf tissues. *EGR2	:GUS* expression was observed in young expanding leaves as well as in recently emerged leaf primordia and was induced by low ψ_w_ in young leaves (Supplemental Fig. S13A and B). However, *EGR2_promoter_:GUS* expression was not detected in the shoot meristem under either condition. Similarly, in *EGR2_promoter_:EGR2-YFP* lines the EGR-YFP fusion protein could not be detected in the shoot meristem (Supplemental Fig. S14). Conversely, *MASP1_promoter_:GUS* expression and native promoter driven MASP1-YFP expression (*MASP1_promoter_:MASP1-YFP*) were detected in the shoot meristem and surrounding leaf primordia (Supplemental Fig. S13 and S14). Low ψ_w_ reduced the level of MASP1 protein in the shoot meristem despite strong MASP1 promoter activity in the shoot meristem at low ψ_w_ (Supplemental Fig. S13 and S14). We could also clearly see *MASP1_promoter_:GUS* expression in the base (close to the petiole) of young leaves where cell division to drive leave blade expansion occurs and could see that in this tissue it overlapped with *EGR2_promoter_:GUS* expression (Supplemental Fig. S13). Thus, separate and overlapping domains of MASP1 and EGR expression may be involved in controlling shoot and leaf growth during low ψ_w_ in a manner similar to their role in controlling root growth.

## Discussion

The rate and duration of cell division and extent of cell expansion are all potential points of control to alter plant growth in response to environmental signals. Which of these control points are most important for modulating growth during abiotic stress and the signaling proteins involved are important questions for plant stress research. Developmental studies have advanced the idea that opposing gradients of regulatory gene expression across the meristem control the transition from cell division to cell expansion (Wendrich et al., 2017; Salvi et al., 2020). Although many such gene expression gradients have been observed, there is scant data to show whether this leads to differences in the stoichiometric ratios of protein abundance at different positions along the root meristem and little data to show how proteins with spatial differences in stoichiometry can influence each other’s activity in a manner consequential for growth regulation. Our results bring these two lines of inquiry together by demonstrating that differing spatial patterns of EGR-MASP1 protein abundance are a key factor regulating root meristem function during low ψ_w_ stress. Disrupting the spatial pattern of EGR-MASP1 protein stoichiometry, either by ectopic expression or by loss of function mutations, changes how root growth responds to low ψ_w_. Under low ψ_w_, phosphorylated MASP1 promotes continued cell division in the proximal root meristem. However, in more distal locations MASP1 protein level decreases and EGRs inactivate remaining MASP1, particularly in the cortex, a cell type important for controlling growth responses to low ψ_w_ (Dietrich et al., 2017). This hastens the transition from cell division to cell expansion (Fig. 5D). Ectopic expression of either EGR2 or MASP1 upsets this balance allowing a longer duration of cell division with higher rates of cell division in the distal meristem region (in the case of *35S:MASP1*) or an earlier exit from cell division and smaller meristem (in the case of *35S:EGR1*). Ectopic expression of MASP1 counteracts the effect of low ψ_w_ to restrict MASP1 expression to a smaller region just behind the QC. Conversely, ectopic expression of EGR1 amplifies the effect of low ψ_w_ to push EGR expression closer to the QC. Thus, the opposing action of MASP1 and EGRs, and differences in the stoichiometric ratio of EGR-MASP1 protein abundance at different positions along the root meristem, allow cell division in the proximal meristem to be protected while also allowing the plant to down-regulate meristem activity and decrease growth during moderate severity low ψ_w_ stress (Fig. 5D). This model of how EGR-MASP1 signaling controls root growth is also supported by previous data demonstrating that the EGR phosphatases interact with MASP1 and that changing the EGR expression level changes the level of MASP1 phosphorylation (Bhaskara et al., 2017). Note that we do not exclude the possibility that EGRs and MASP1 also affect the rate of cell division (cell cycle time) within the meristem since *egr1-1egr2-1* had higher cell division activity at low ψ_w_ while having similar meristem size as wild type and *35S:MASP1* had a larger effect on meristem cell number than on meristem length.

*35S:EGR1* did not decrease growth, cell division, cell size or meristem size of unstressed plants despite having strong effect on plants exposed to low ψ_w_ and despite similar levels of root tip *35S*-driven EGR1 protein expression in both control and stress treatments. One possible explanation is that EGR phosphatase activity is inhibited (or not activated) in the root meristem of unstressed plants. Such post-translational regulation of EGR activity could occur via interaction with regulatory proteins in a manner similar to inhibition of Clade D PP2Cs by SAUR proteins (Spartz et al., 2014), which is also important for growth regulation, or similar to PYL/RCAR inhibition of Clade A PP2C activity which is of central importance to ABA signaling (Cutler et al., 2010; Raghavendra et al., 2010). As we have previously discussed, it is possible that additional PP2Cs, including Clade E PP2Cs, are paired with regulatory proteins that control their activity (Bhaskara et al., 2019). However, we cannot rule out other explanations such as the possibility that EGRs are active in unstressed plants but their effect masked by other mechanisms that control meristem activity, the phosphatase is active but access to its substrate proteins (including MASP1) is restricted in unstressed plants or, EGR substrate proteins only become phosphorylated in stress-treated plants. We also note that high expression of EGR1 in cell types other than the cortex could be a factor in the dramatic effect of *35S:EGR1* on root meristem size during low ψ_w_ stress. However, the observation that *egr1-1egr2-1* had increased cell division at low ψ_w_, while *egr1-1egr2-1masp1-1* did not, indicates that EGR regulation of MASP1 phosphorylation status in cortical cells, where EGR2 is most highly expressed, can be sufficient to affect cell division activity of the meristem as a whole. None of our mutant or ectopic expression lines, had altered root meristem morphology in terms of organization of cell files and placement of cell division planes. How expression of EGR2 predominantly in the cortical cell layer affects root cell division and cell expansion during low ψ_w_ stress in way that preserves the coordination between different cell layers is an interesting question for future research.

EGR2 protein level was dramatically higher in cortex compared to other root cell types. This is consistent with recent root tip single-cell RNA sequencing which found EGR expression in most cells that clustered with the cortex and some cells in the epidermal cluster (Wendrich et al., 2020). The same data set showed MASP1 expression across all root cell types. The cortex-enriched expression of EGR2 is particularly interesting in light of recent observations that root cortex is the key cell layer for response to a ψ_w_ gradient during hydrotropic bending (Dietrich et al., 2017). Dietrich et al. (2017) demonstrated that expression of SnRK2.2/SnRK2.3 in cortical cells was critical for differential cell elongation during hydrotropic root bending. Interestingly, EGR2 has been shown to inhibit the kinase activity of SnRK2.6 (also known as Open Stomata 1, OST1; Ding et al., 2019) which is closely related to SnRK2.2 and SnRK2.3. Thus, it will be of interest to investigate whether EGR regulation of SnRK2 activity in the root cortex is one factor that limits cell elongation during low ψ_w_ or is involved hydrotropic root bending. Another recent study has also proposed that inter-cellular signaling originating from the cortex is important for controlling root growth in response to environmental signals (Mielke et al., 2021). It is similarly interesting to note that altered hydrotropic response 1 (*ahr1*), which is putatively involved in hydrotropic root bending, also has increased root elongation and maintains larger root meristem size than wild type during low ψ_w_ stress (Salazar-Blas et al., 2017). Unfortunately, the mutated gene responsible for the *ahr1* phenotype has not yet been identified.

It is also worth noting that *egr1-1egr2-1* did not have detectable effect on root cell division of unstressed plants but did affect cell size in both control and stress treatments. This again is consistent with the idea that EGRs affects cell size via a different, MASP1-independent, mechanism and suggests that EGR activity may also be regulated differently during cell expansion and maturation than in the meristem. We also observed that, *35S:MASP1* had a similar increased cell size as *egr1-1egr2-1* and *egr1-1egr2-1masp1-1*. While it is tempting to speculate that this is related to MASP1 effects on microtubule stability (Bhaskara et al., 2017), a perhaps simpler explanation is that ectopic MASP1 expression in the cell expansion zone, where endogenous MASP1 is very low, titrates away EGR phosphatase activity and thus allows greater cell expansion. Our data confirmed that MASP1 S670 phosphorylation is critical for its function. Thus, kinases that phosphorylate MASP1 S670 are also potential regulators of meristem size and function.

Our observations that low ψ_w_ decreases root cell division; meristem size and cell expansion are similar to observations of West et al. (2004) who studied the basis for decreased root elongation in wild type Arabidopsis exposed to mild salt stress. In their case, salt stress reduced meristem size and cell production (measured both kinematically and using the CYCB1::GUS reporter) and also reduced final cell length. The increased growth and increased root meristem size of *egr1-1egr2-1* and *35S:MASP1* were also similar in some ways to the recently reported effects of β-cyclocitral which increased root meristem size (but did not affect cell length) when applied to unstressed plants (Dickinson et al., 2019). β-cyclocitral also stimulated growth during salt stress; although, whether this was due to increased meristem size and whether β-cyclocitral affects growth during drought stress are yet to be reported. Nonetheless, a common theme that emerges from these results is that, whether by genetic or chemical means, inducing plants to maintain a larger population of dividing cells (larger meristem) can allow enhanced growth maintenance after exposure to abiotic stress.

## Materials and Methods

### Plant materials and stress treatment

The *egr1-1egr2-1*, *masp1-1* and *egr1-1egr2-1masp1-1* mutants have been previously described (Bhaskara et al., 2017). The meristem analysis used previously described *35S:YFP-MASP1* line as this construct could complement the *masp1-1* mutant and was functionally equivalent to *35S:MASP1* (untagged MASP1) in our previous analysis (Bhaskara et al., 2017). Data presented in Fig. 1-3 use both *35S:MASP1* and *35S:YFP-MASP1* lines as well as *35S:EGR1* and *35S:YFP-EGR1* to ensure that multiple independent transgenic lines were used for each construct and to ensure that lines expressing the YFP-tagged proteins had the same phenotypic effects as lines expressing untagged protein. Details of the construction of these lines and vectors used are given in Bhaskara et al. (2017). To analyze the effects of MASP1 phosphorylation, the *35S:MASP1^S670A^/GFP-TUA6* and *35S:MASP1^S670D^/GFP-TUA6* lines described by Bhaskara et al. (2017) were used. It had been previously demonstrated that that the presence of the GFP-TUA6 marker did not affect root elongation under either control or stress treatments (Bhaskara et al. 2017). For clarity of presentation, these lines are referred to simply as *35S:MASP1* and *35S:EGR1* in figures and text. The CYCB1::GUS reporter lines were generated by crossing the mutants, *35S:YFP-MASP1*, *35S:YFP-EGR1* and 35S:EGR1 lines to the CYCB1:GUS reporter line and F_3_ homozygous lines were used for cell division assays. Additionally, the CYCB1:GUS reporter line was transformed with untagged MASP1 (in pEG100 vector) and both lines used for the root elongation and cell division assays in Fig. 1 and 2. In all cases, lines with a single locus insertion of the transgene (based on segregation ratio of the selectable marker in the T_2_ generation) were isolated and T_3_ homozygous seed stocks used for experiments of for making crosses. Two to four independent transgenic lines were used for all experiments. As the data were indistinguishable between transgenic lines, combined data are shown in the main text figures (Fig. 1, 2, 3) while data for individual transgenic lines is shown in Supplemental Figures S2, S4, and S7). A list of all transgenic lines used in this study, and the experiments they were used for, is shown in Supplemental Table S1.

For analysis of native promoter driven MASP1 and EGR2 protein levels and expression pattern, C-terminal YFP constructs (*MASP1_promoter_:MASP1-YFP* and *EGR2_promoter_:EGR2-YFP*) were generated by amplifying the promoter and coding region of each gene (3663 bp and 3671 bp for *MASP1* and *EGR2,* respectively) from genomic DNA of Col-0. The PCR products were first cloned in entry vector (pDNOR221) and were then transferred to expression vector (pGWB540) via Gateway recombination (primer sequences are shown in Supplemental Table 1). The constructs were subsequently transformed into *masp1-1* and *egr1-1,2-1* mutants (and Col-0 wild type in the case of *EGR2_promoter_:EGR2-YFP*) through *Agrobacterium*-mediated floral dip method. *Promoter:GUS* constructs were generated by cloning the promoter region (1347 bp and 1653 bp upstream of start codon for *MASP1* and *EGR2,* respectively, primer sequences used for cloning are shown in Supplemental Table S2) into pGWB443 by Gateway cloning and transformation into Col-0 wild-type. Homozygous T_3_ seed stocks of lines having a single locus transgene insertion were isolated as described above. Two or three independent transgenic lines were used for subsequent experiments. Lines with *MASP1_promoter_:MASP1-YFP* in the wild type background were generated by crossing *MASP1_promoter_:MASP1-YFP/masp1-1* to wild type to segregate the *masp1-1* T-DNA insertion away from the *MASP1_promoter_:MASP1-YFP* transgene.

Seedling growth and low ψ_w_ treatment on PEG-infused agar plates was performed as described previously (Verslues et al., 2006; Bhaskara et al., 2017). For seedling growth assays, seeds were plated on unstressed control media, stratified for three days and transferred to a growth chamber (22 °C, continuous light intensity of 100-120 μmol m^-2^ s^-1^). Five-day-old seedlings were transferred to fresh half-strength MS plates (control, −0.25 MPa) or plates infused with PEG-8000 (−0.7 MPa). Plates were always prepared and infused with PEG 15 h before use to avoid drying of the media and generate plates of consistent ψ_w_. All media was prepared without addition of sugar. Primary root elongation rates were measured by marking the position of the root apex on the back of the plate at 48 h intervals for six days after transfer. Root elongation rates were combined from two or three independent experiments.

Other experiments quantifying total root elongation and seedling fresh or dry weight (presented in Supplemental Fig. S10 and S11) were conducted as described in Bhaskara et al. (2017). Five-day-old seedlings were transferred to fresh control plates or −0.7 MPa PEG-infused plates. In the control treatment, root elongation was measured over the subsequent five days (when the seedlings nearly reached the maximum size accommodated by the agar plates) and seedling fresh and dry weights measured at the end of the experiment. The same procedure was followed for the stress treatment except that the seedlings were allowed to grow for ten days after transfer to reach a developmental stage close to that obtained by unstressed seedlings at five days after transfer. For these experiments, growth of the transgenic plants was normalized versus wild type seedlings growing in the same plate. Three or four independent experiments were performed with each experiment containing two plates for each genotype analyzed. The experiment averages were tested for significant differences from wild type using one sample T-tests.

### GUS, EdU and PI staining

β-Glucuronidase (GUS) activity staining for *promoter:GUS* and CYCB1::GUS lines were performed using standard procedures. Seedlings were immersed in ice cold 90 % acetone for 10 min, washed once with water and then put into staining solution (50 mM phosphate buffer pH 7.2, 100 μM K_3_Fe(CN)_6_, 100 μM K_4_Fe(CN)_6_, 1 mM X-GLUC in dimethyl formamide) and vacuum infiltrated for 20 min followed by incubation at 37 °C for 2 to 5 h. The samples were then transferred to 80% ethanol and keep at 4 °C for overnight. Next day, the samples were cleansed with a solution of methanol and acetic acid (3:1) for 5-10 min and then transferred to 80% ethanol before taking images (Zeis Imager Z1). Whole rosette images (Supplemental Fig. S13) were obtained using the tiles function of the Zeiss Zen software package.

For 5-ethynyl-2′-deoxyuridine (EdU) staining, seedlings were grown on 1/2 MS agar plates for 5 days and transferred to control or stress plates (−0.7 MPa) for four days. The seedlings were labelled by pouring EdU (Thermo Fisher Scientific) solution (10 μM EdU in ½ MS or ½ MS + −0.7 MPa PEG) into the plates and incubating for 15-30 min in a growth chamber. Roots were transferred to a 2 mL tube and fixed for 20 min in 4% (w/v) paraformaldehyde in PBS (pH 7.4), washed twice with PBS (2×10 min) and placed in 0.5% (v/v) TritonX-100 in PBS. After 20 min, samples were washed with PBS twice and incubated in 500 μL of Click-iT reaction cocktail (1× PBS, 4 mM CuSO4, 40 mM sodium ascorbate and 5 μM AF594 azide) for 30 min in the dark. The Click-iT reaction cocktail was removed and samples were washed with PBS once before observing under confocal microscope. Fluorescence spots within the apical 300 μm of the root tip were counted.

For Propidium Iodide (PI) staining, seedlings were immersed in PI solution (0.01 mg/mL of PI in water or 0.01 mg/mL of PI in 285 mM mannitol (to match the water potential of the −0.7 MPa stress and prevent cell swelling) for 60 sec (control) or 90 sec (stress) and rinsed with water/mannitol solution before mounting on glass slide for confocal microscopy. For meristem size analysis of seedlings 4 days after transfer to stress or control plates (Supplemental Fig. S4) or younger 2 or 4 day-old seedlings (Supplemental Fig. S5), roots were cleared with a 8:3:1 mixture of chloral hydrate:water:glycerol, mounted on a glass slide and meristem cells counted as previously described (Biancucci et al., 2015)

### Confocal microscopy and image analysis

Confocal laser-scanning microscopy was done using a Zeiss LSM510 for quantification of meristem cell numbers and meristem size as well as EdU staining. For other high resolution imaging a Zeiss LSM880 or Leica Stellaris 8 (Fig. 4C images) was used. Low ψ_w_-treated seedlings were mounted in solution of the same ψ_w_ (−0.7 MPa) as the stress treatment to prevent swelling during imaging. EdU or PI stained samples were visualized using excitation at 543 nm and emission at 588nm while YFP imaging used excitation at 514 nm and emission at 542 nm. To quantify the protein level of YFP-tagged EGR or MASP1 across the root tip (Fig. 4B, Supplemental Fig. S12), roots were imaged with a fluorescence imager (Zeiss Axio Imager Z1) and 2 or 3 images covering the apical 1 cm of the root merged using the tiles function of Zeiss Zen software and analyzed using the line scan function of Image J. For each root tip, scans of three different lines starting the root tip and extending for approximately 800 μm of the root were averaged. Scans followed the area between the stele and epidermis of the root where fluorescence intensity was highest. Line scans of the area alongside the root were used for background subtraction. The data presented in Fig. 4B, and Supplemental Fig. S12A and B, are means of 5-10 roots per genotype and treatment combined from two independent transgenic lines. For higher resolution visualization of the whole root tip, merged images of the root tip were obtained by using the tiles function of either Zeiss ZEN software (Supplemental Fig. S10 and S11) or Leica LAS X software (Fig. 4C). Radial images of root were obtained by collecting a Z-stack of root tip images and processing using the Fiji orthogonal viewer. To image the shoot meristem, seedlings were first dissected under a stereo microscope and the meristem region mounted and imaged with a Zeiss LSM880 microscope.

## Acknowledgements

We thank Arnould Savoure (UPMC, Sorbonne Universités) for providing seed of the CYCB1::GUS reporter line, the staff of the imaging core facility of the Institute of Plant and Microbial Biology (Ji-Ying Huang and Mei-Jane Fang) for microscopy assistance and Shih-Shan Huang for laboratory assistance. This research was supported by an Academia Sinica Investigator Award (AS-IA-108-L04) and Taiwan Ministry of Science and Technology grant (MOST 107-2311-B-001-037) to P.E.V.

## Author contributions

P.E.V. conceived research, T.L., G.B.B and P.E.V. designed experiments, T.L., C.Y.C, M.B. and G.B.B. performed experiments, T.L., C.-Y.C. and P.E.V. analyzed data, P.E.V. wrote the manuscript with assistance from T.L. and G.B.B. All authors read and approved the manuscript.

---

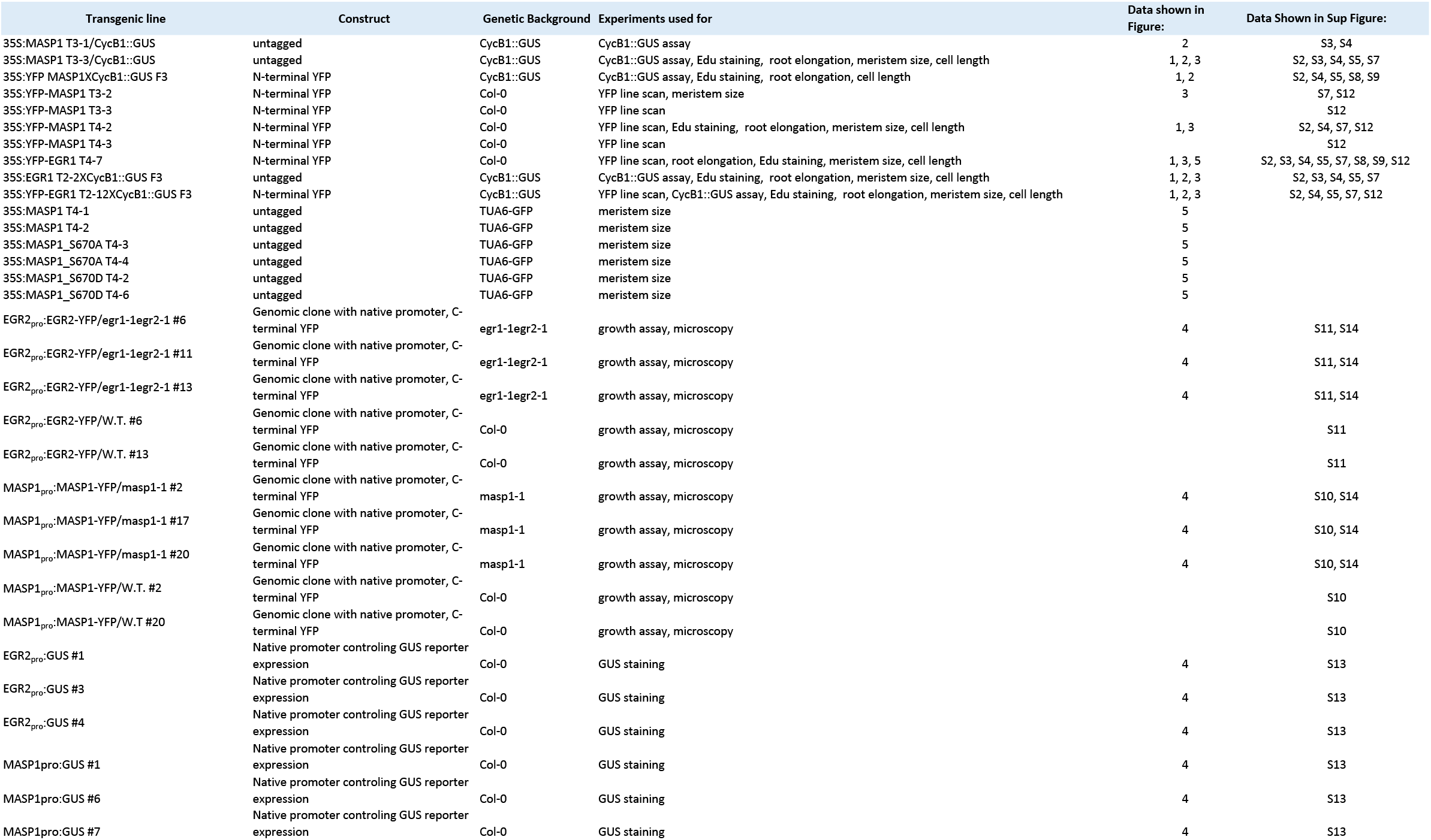

