## Supplemental Figures 1-14 for "Spatial differences in stoichiometry of EGR phosphatase and Microtubule-Associated Stress Protein 1 control root meristem activity during drought stress"

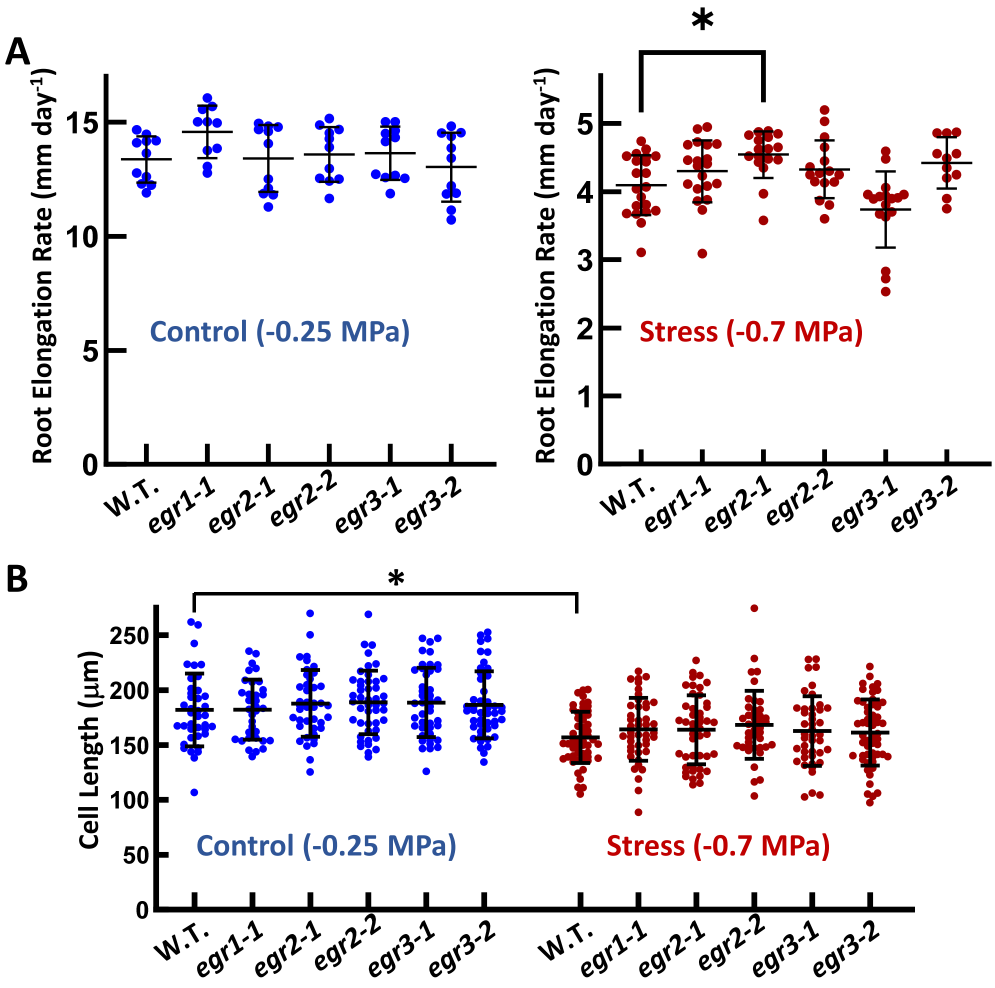


**Supplemental Figure S1: Root Elongation and cell length of *egr* single mutants (Supports Fig. 1 and 2).**

A. Five-day-old seedling of each genotype were transferred to either fresh control plates or -0.7 MPa agar plates for stress treatment. Root elongation was measured over the subsequent six days. There were no significant differences between wild type and *egr* mutants in the unstressed control. In the stress treatment, *egr2-1* has significantly higher (ANOVA with Tukey's post hoc test test, corrected $P \leq 0.05$) root elongation rate than wild type (W.T.). Data are combined from two independent experiments ($n = 11-20$) . Error bars indicate the standard deviation.

B. Epidermal cell lengths measured at 2 mm from the root apex at 4 days after transfer of five-day-old seedlings to control or stress (-0.7 MPa) plates. Stress significantly decreased cell length of W.T. (T-test, $P \leq 0.001$) but there were no significant differences between wild type and mutants within the control or stress treatments (ANOVA with Tukey's post hoc test test, corrected $P \leq 0.05$). Data are combined from two independent experiments ($n = 34-56$). Error bars indicate the standard deviation .

The *egr* T-DNA mutants have been previously described in Bhaskara et al. (2017).

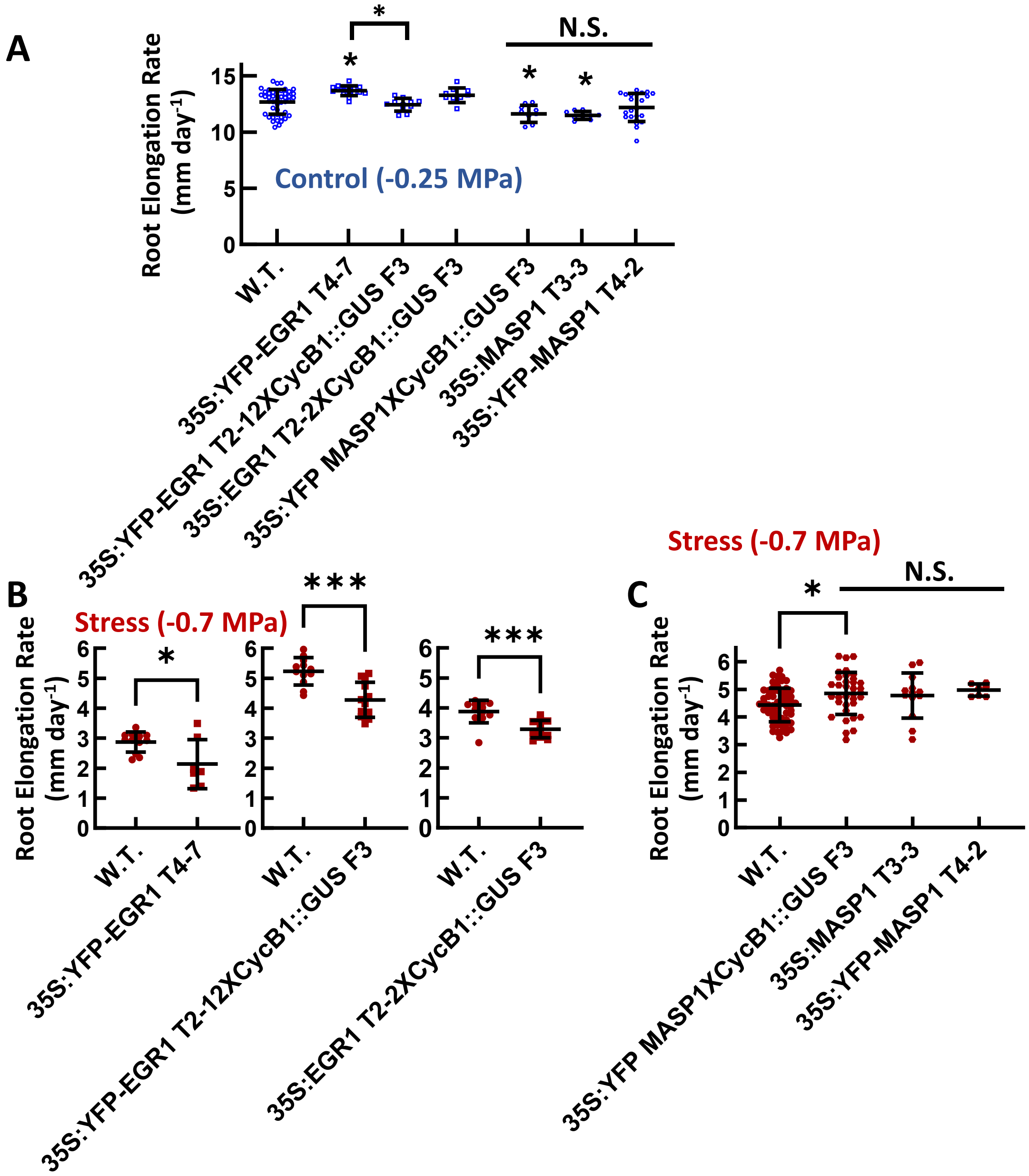


**Supplemental Figure S2: Additional root elongation data for individual *35S:EGR1* and *35S:MASP1* transgenic lines (Supports Fig. 1).**

Five-day-old seedlings of wild type and 35S lines were transferred to fresh control plates (A) or low water potential (-0.7 MPa) plates (B and C) and root elongation measured over the subsequent six days. In A, data are combined from two independent experiments (N = 10-23 for 35S-lines , 46 for W.T.). In B, data are from single experiments that are representative of other replicate experiments (N = 7-12). In C, data are combined from 2 independent experiments except for T4-2 which is from one experiment (N = 7-63).

Asterisks directly above the data points in A indicate significant difference compared to wild type. Other comparisons among EGR1 or MASP1 transgenic lines are indicated by brackets and bars above the data. Significant differences compared to wild type based on ANOVA (A and C) or T-test (B). Error bars indicate standard deviation. N.S. = Non Significant.

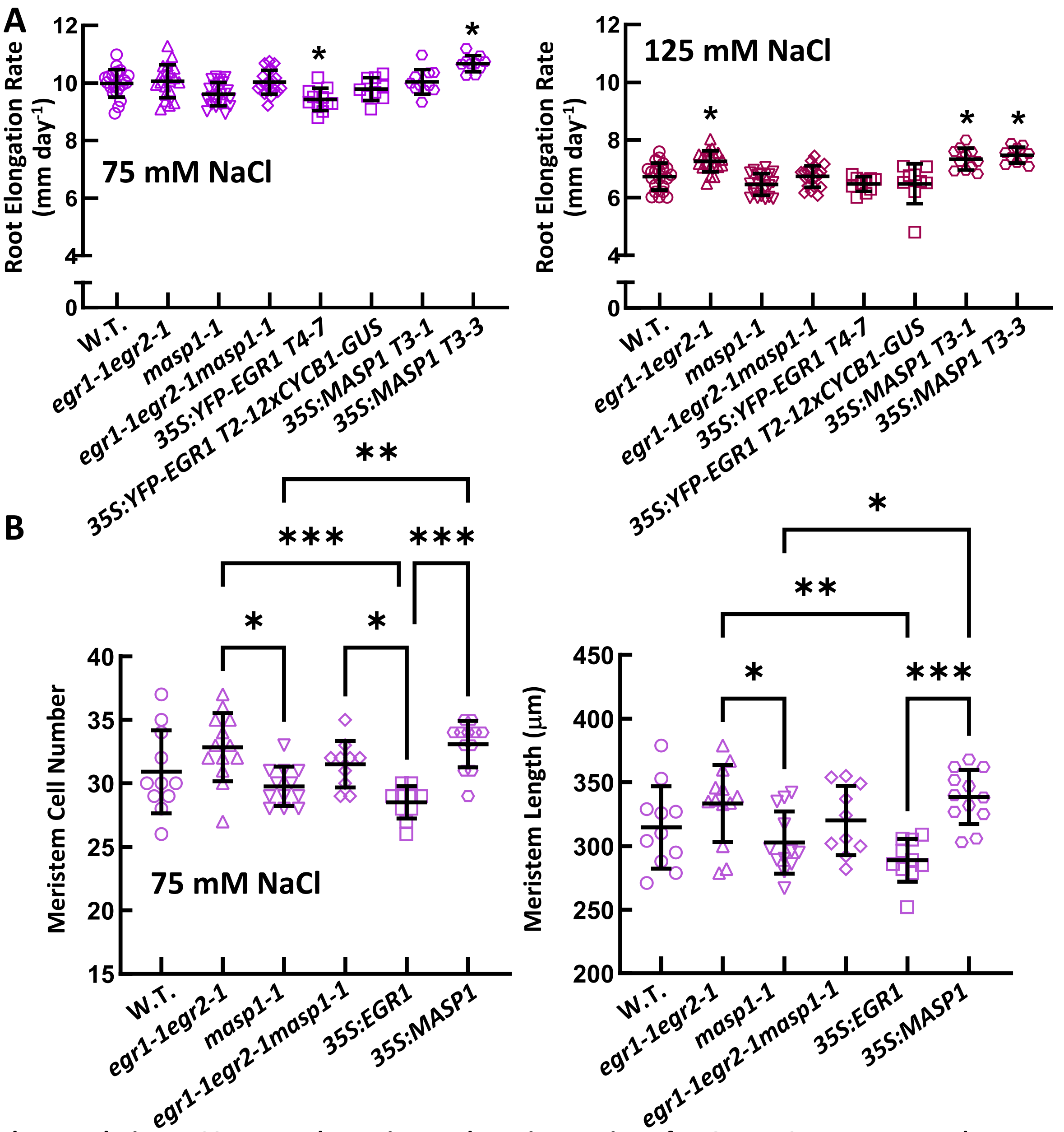


**Supplemental Figure S3: Root Elongation and meristem size of EGR-MASP1 mutant and transgenic lines exposed to moderate severity salt stress (Supports Fig. 1 and 3).**

A. Five-day-old seedling of each genotype were transferred to agar plates containing the indicated NaCl concentrations and root elongation measured over the subsequent six days. For 75 mM NaCl, none of mutant or transgenic lines were significantly different than W.T. but did differ from each other as indicated. Asterisks indicate significant differences compared to W.T. (ANOVA with Tukey's post hoc test, corrected $P \leq 0.05$; n = 20-21). Error bars show standard deviation (S.D.).

B. Root meristem cell number and length measured five days after transfer to 75 mM NaCl. For *35S:EGR1* and *35S:MASP1*, data are combined from two independent transgenic lines. Brackets and asterisk (*) indicate significant differences between genotypes (ANOVA with Tukey's post hoc test test, corrected P ; *, $P \leq 0.05$; **, $P \leq 0.01$; ***, $P \leq 0.001$; n = 11-13). Error bars show S.D

For both A and B, data are combined from 2 or 3 independent experiments.

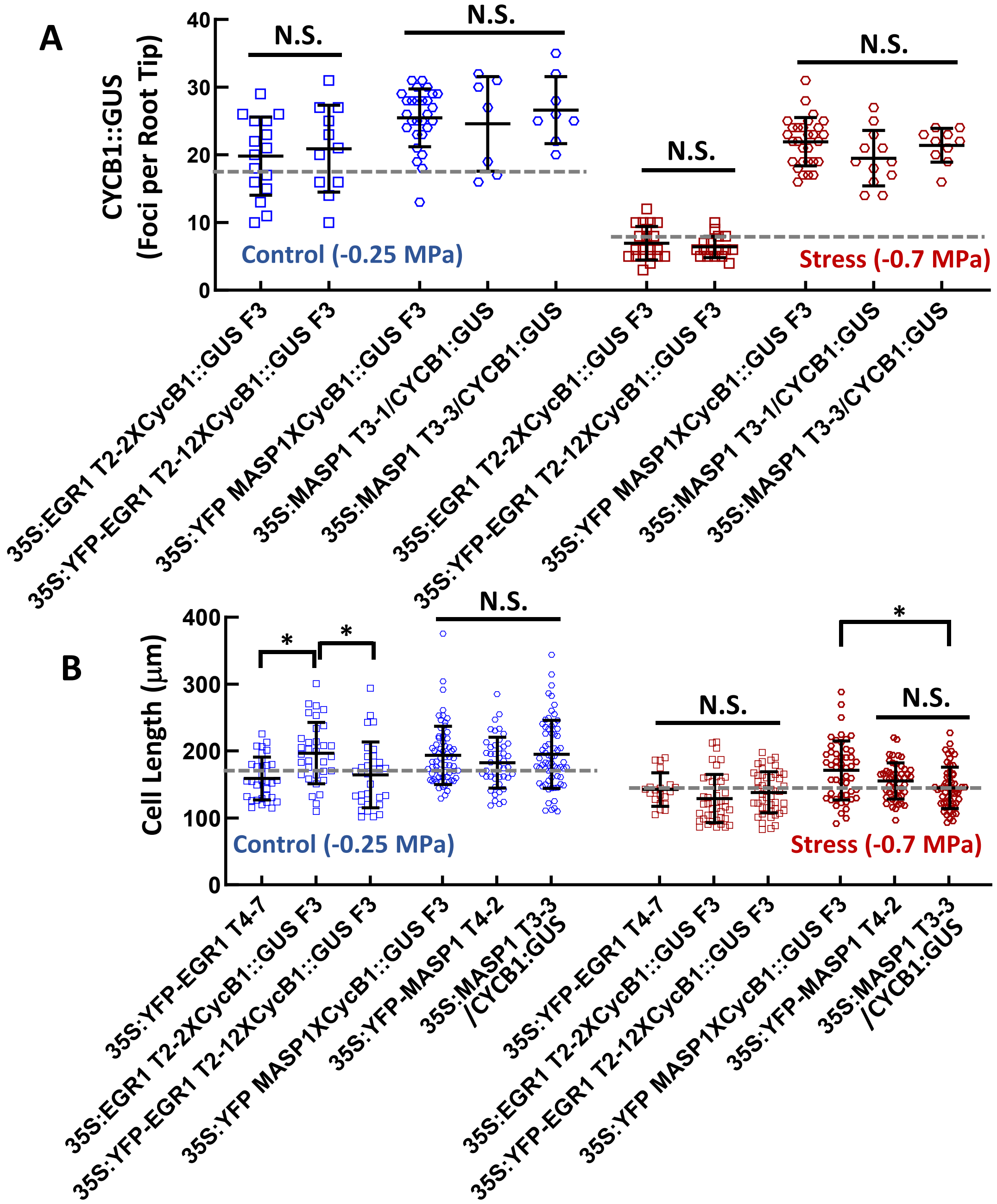


**Supplemental Figure S4: CYCB1::GUS foci counts and epidermal cell lengths for individual EGR1 and MASP1 transgenic lines. (Supports Fig. 2).**

A. Quantification of CYCB1::GUS foci in the primary root meristem of individual transgenic lines measured 4 days after transfer to the indicated treatments. Symbols indicate counts of GUS foci from individual roots while black bars and error bars indicate the mean and standard deviation for each genotype. Data are combined from two independent experiments for each line (n = 10-28)

B. Cell length at 2 mm from the root tip measured four days after transfer of mutant or ectopic expression seedlings to control or stress treatment. N = 20-66

For A and B, significant differences among EGR1 or MASP1 transgenic lines (ANOVA, P ≤ 0.05) are indicated by brackets and asterisks. N.S. = Non Significant. Any comparison among EGR1 transgenic lines or among MASP1 transgenic lines in control or stress treatments not specifically marked is also non significant. Dashed gray lines indicate the mean value of wild type (Fig. 2)

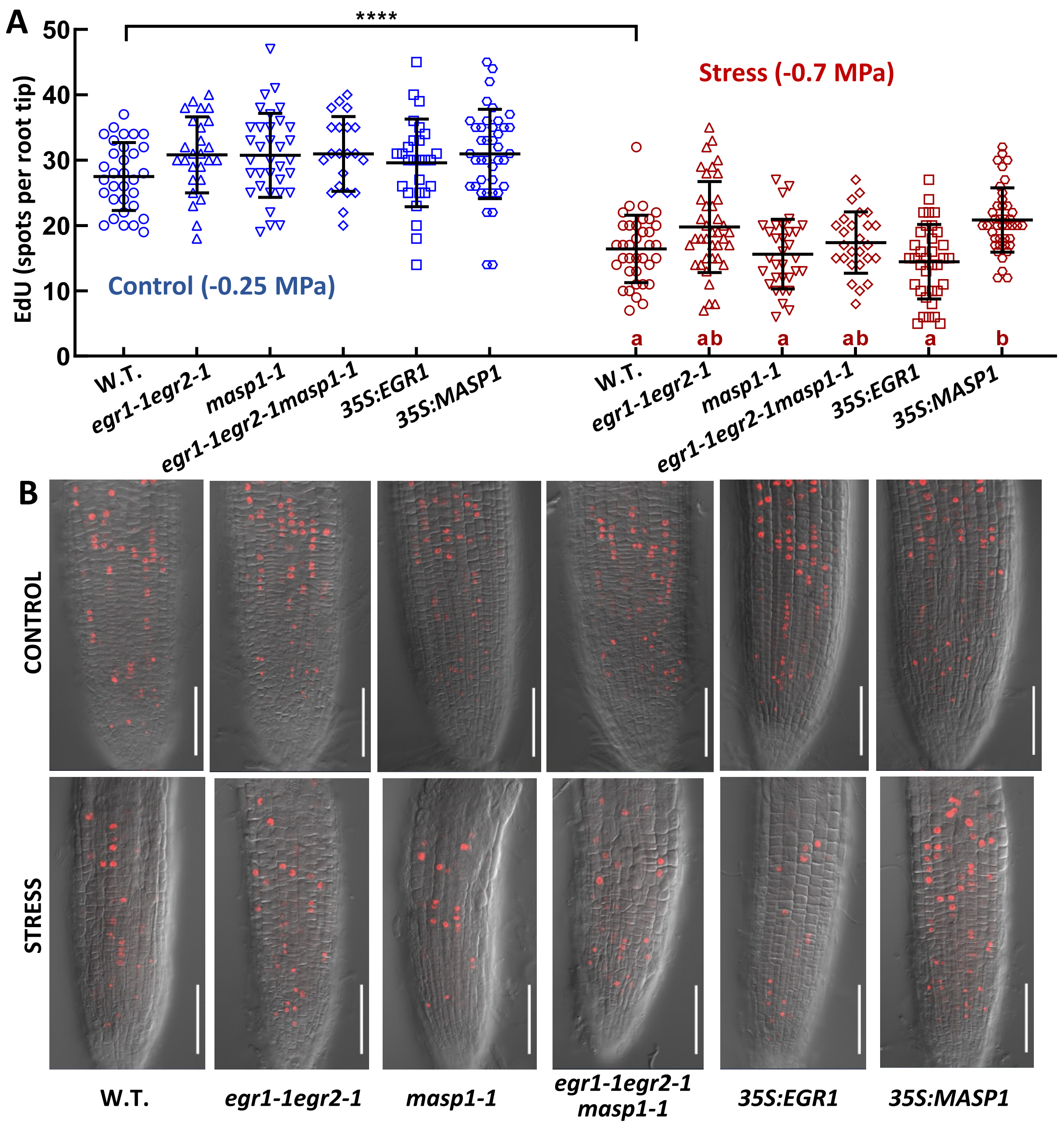


**Supplemental Figure S5: EdU staining confirms that EGRs and MASP1 affect cell division during low $\psi_w$ stress (Supports Fig. 2).** Five-day-old seedlings were transferred to control or low $\psi_w$ (-0.7 MPa) for 4 days and then used for EdU staining (10 µM EdU; 15 minute incubation for unstressed control, 20 minute incubation for stress treated seedlings).

A. Quantitation of EdU foci in the apical 300 µm of the primary root tip. Data are combined from three independent experiments. Within the control or stress treatments, genotypes sharing the same letter do not significantly differ from one another (ANOVA with Tukey's post hoc test, corrected P ≤ 0.05, error bars indicate S.D.). No significant differences between genotypes were detected in the unstressed control. Wild type did significantly differ between control and stress treatment (T-test, P ≤ 0.001). Data for *35S:EGR1* and *35S:MASP1* are combined from multiple independent transgenic lines (n = 21-40).

B. Representative images of EDU stained root tips. Scale bars = 100 µm. Note that cells of *masp1-1*, and to lesser extent *egr1-1egr2-1masp1-1*, are more prone to anisotrophic swelling (Bhaskara et al., 2017) and thus became distorted during the EDU staining process. This did not affect the counting of EDU foci or interpretation of data.

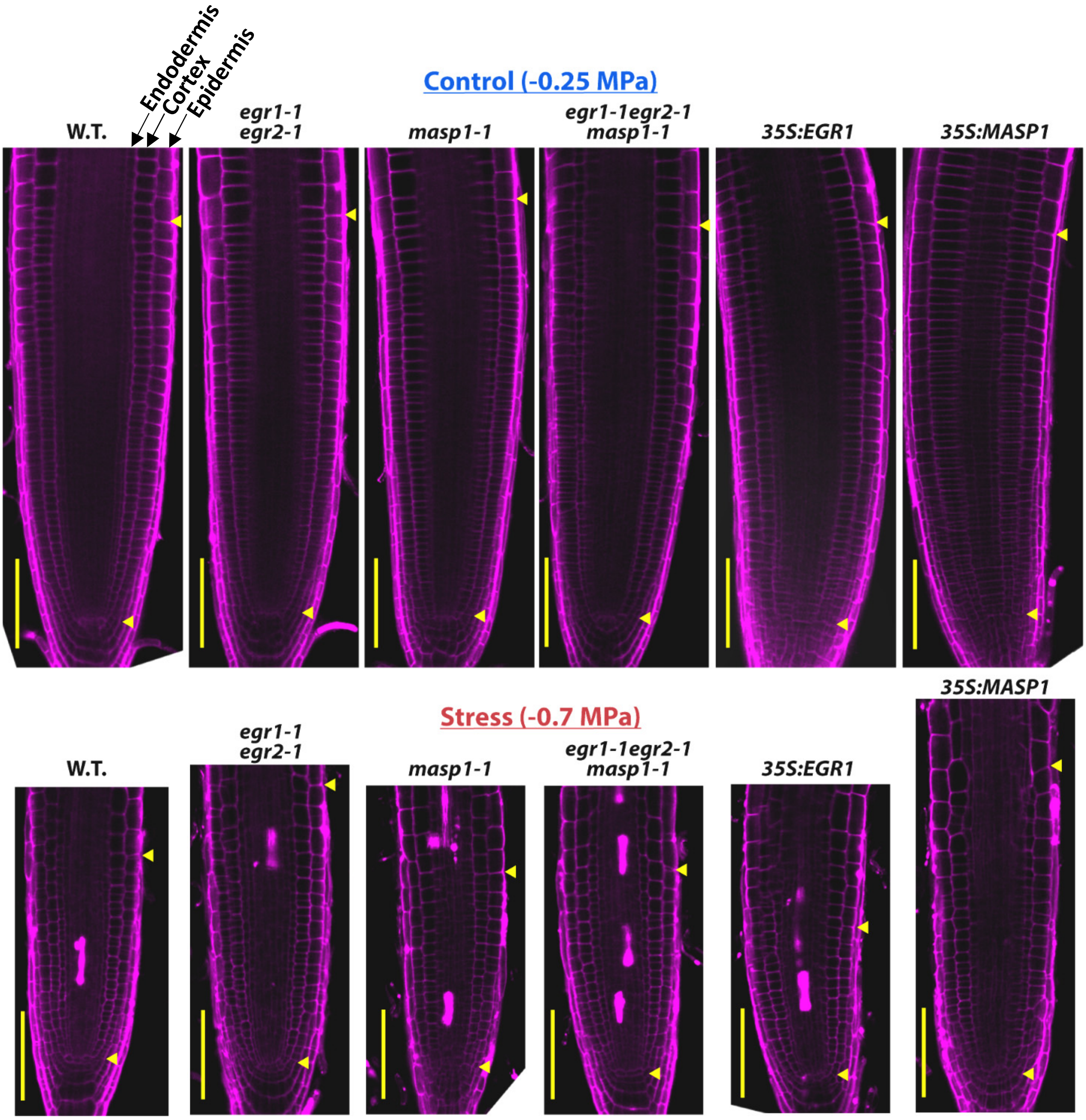


**Supplemental Figure S6 (Supports Fig. 3): Representative images of PI stained root tips used for meristem cell size measurements in control and stress treatments (Supports Fig. 3B).** Arrows mark the ends of the root meristem. Scale bars indicate 100 μm. The epidermis, cortex and endodermis cell layers are marked on the wild type control root for reference.

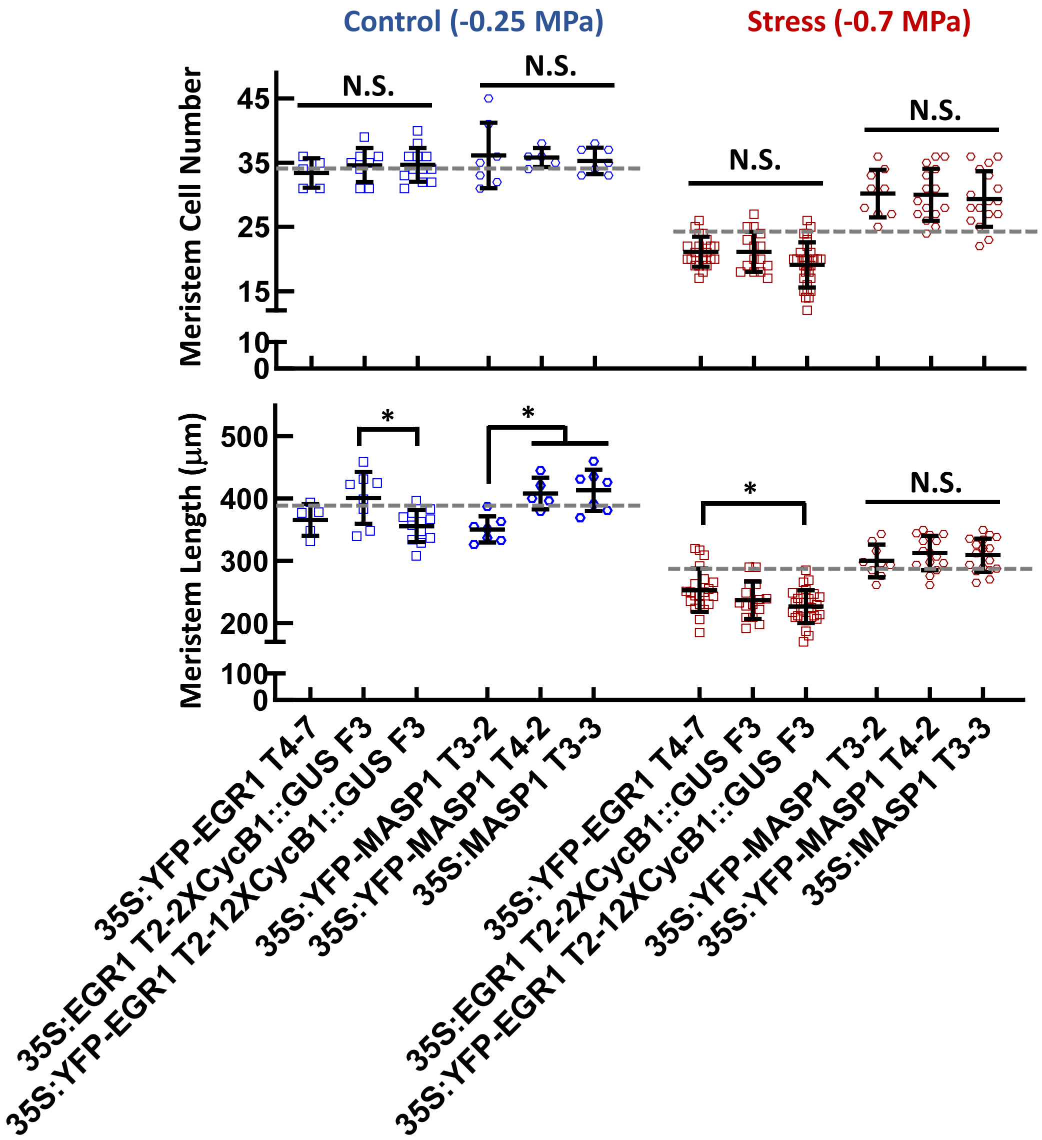


**Supplemental Figure S7: Root meristem cell number and meristem length for individual transgenic lines (Supports Fig. 3B).**

Meristem cell number and meristem length five days after transfer of five-day-old mutant and ectopic expression seedlings to unstressed control or to low $\psi_w$ (-0.7 MPa). Data are combined from 2-3 independent experiments for each genotype. Symbols indicate data from individual roots while black bars and error bars indicate the mean and standard deviation for each genotype (n = 7-29). Significant differences among EGR1 or MASP1 transgenic lines (ANOVA, P ≤ 0.05) are indicated by brackets and asterisks. N.S. = Non Significant. Any comparison among EGR1 transgenic lines or among MASP1 transgenic lines in control or stress treatments not specifically marked is also non significant. Dashed gray lines indicate the mean value of wild type (Fig. 3B)

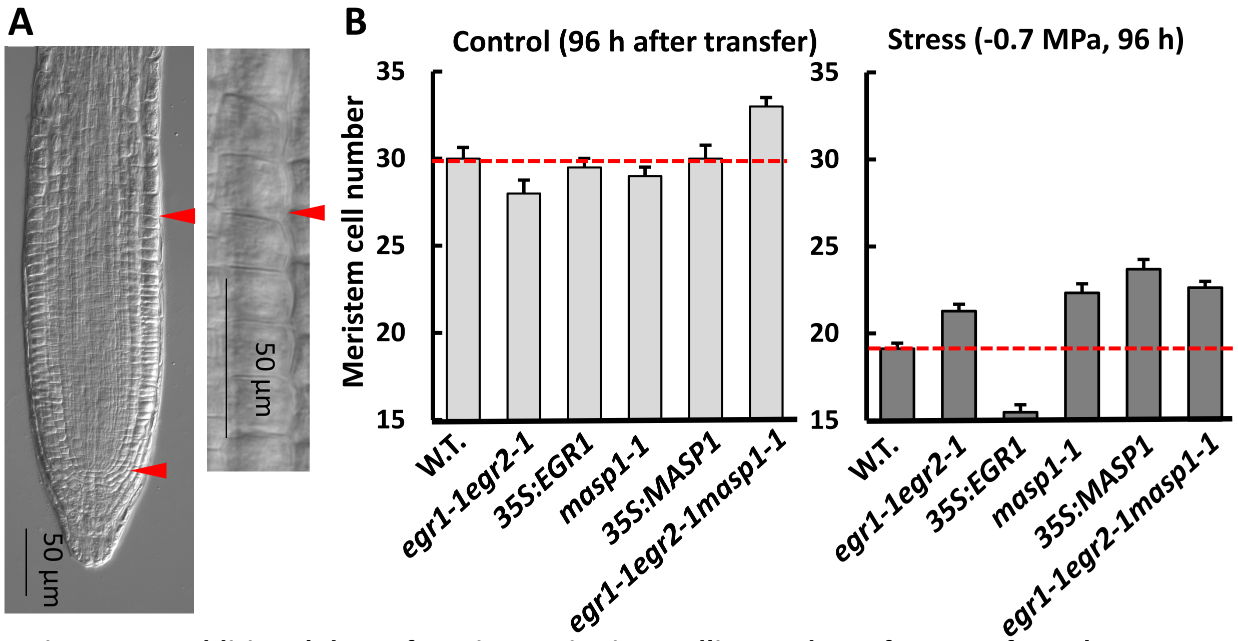


**Figure S8: Additional data of meristem size in seedlings 4 days after transfer to low water potential stress (-0.7 MPa) or control treatments (Supports Fig. 3).**

A. Representative image of wild type primary root tip imaged after chloral hydrate clearing. Red arrows mark the quiescent center and end of meristem.

B. Additional data of meristem size in seedlings 4 days after transfer to low ψw stress (-0.7 MPa) or control. Data are means ± S.E. (n = 10-12)

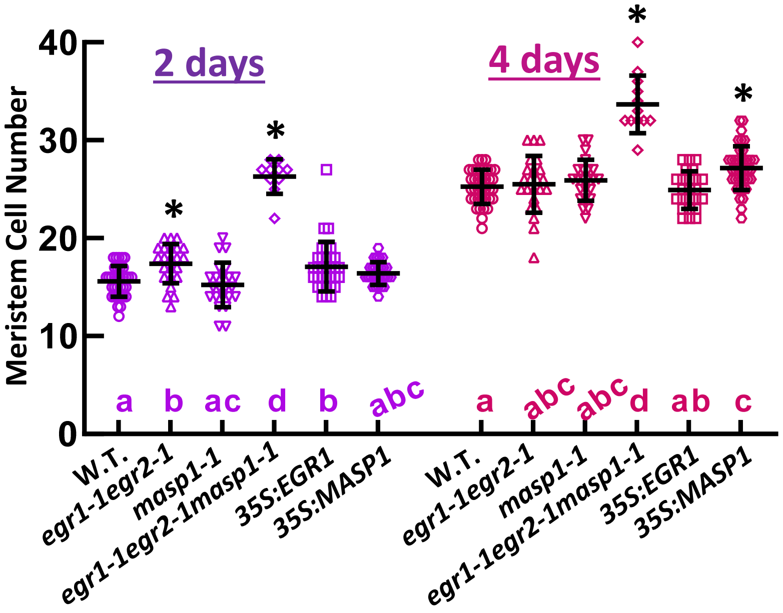


**Supplemental Figure S9: Root meristem cell number in early seedling development (Supports Fig. 3).**

The number of root meristem cells was counted 2 or 4 days after end of the stratification period for seeds plated on unstressed control media**.** Asterisks indicate a significant difference compared to wild type (ANOVA, corrected P ≤ 0.05). Error bars indicate standard deviation. Data are combined from two independent experiments (n = 10-39).

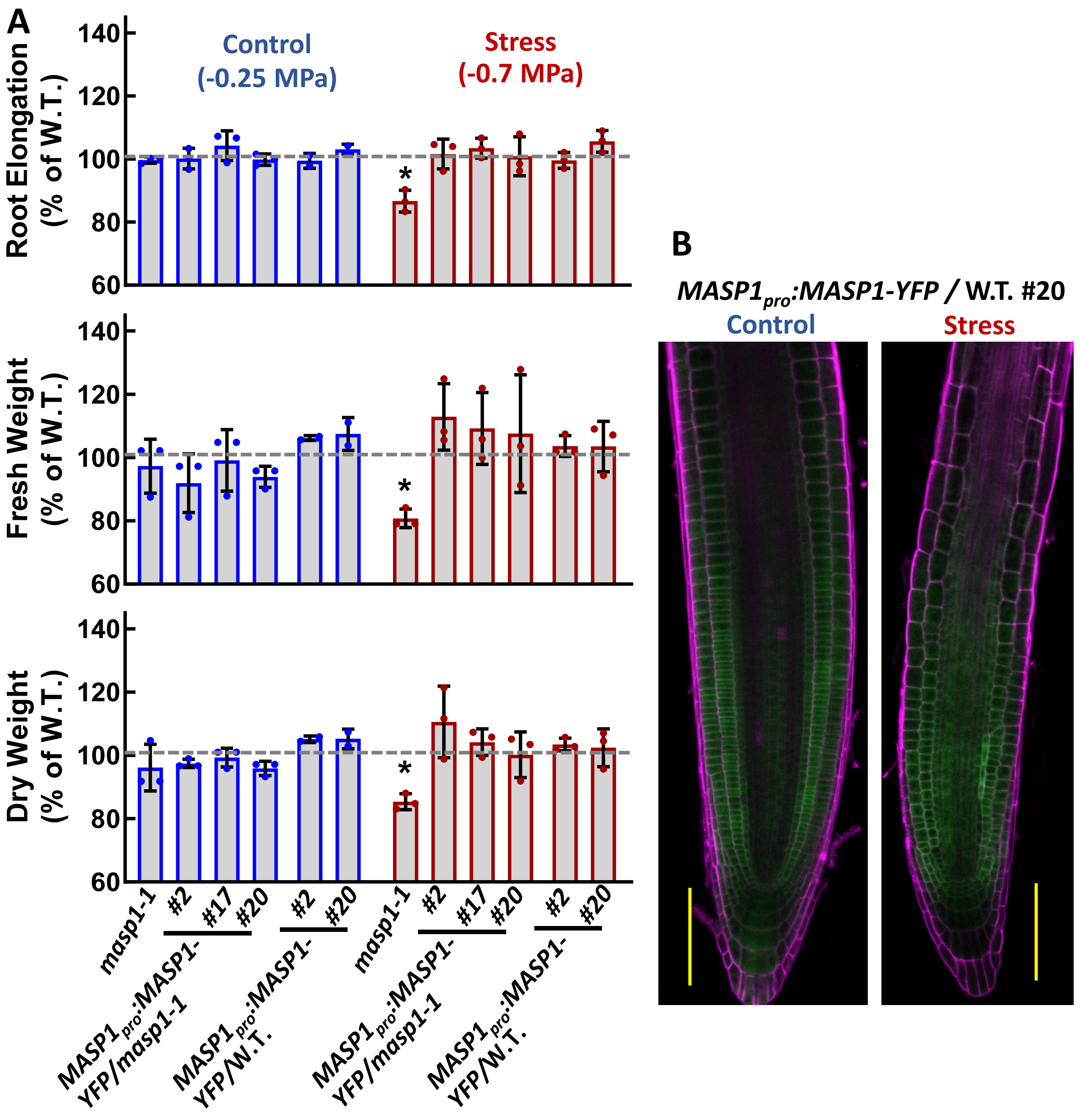


**Supplemental Figure S10: Complementation of *masp1-1* by *MASP1$_{pro}$:MASP1-YFP* (Supports Fig 4).**

Seedling primary root elongation, fresh weight and dry weight were assayed in unstressed control or low $\psi_w$ stress (-0.7 MPa) for *masp1-1,* three independent complementation lines and two lines of native promoter driven MASP1-YFP in the wild type background. Experiments were performed following previously described methods (Bhaskara et al., 2017). Asterisks indicate significant difference from wild type (100%, indicated by the dashed line) by one-sample T-test ($p \leq 0.05$). Data are combined from three independent experiments (n = 3).

B Representative root tip images of the *MASP1$_{pro}$:MASP1-YFP/W.T. #20* transgenic line shown in panel B. YFP fluorescence is shown in green, PI staining of cell wall is shown in magenta. Scale bars indicate 100 μm.

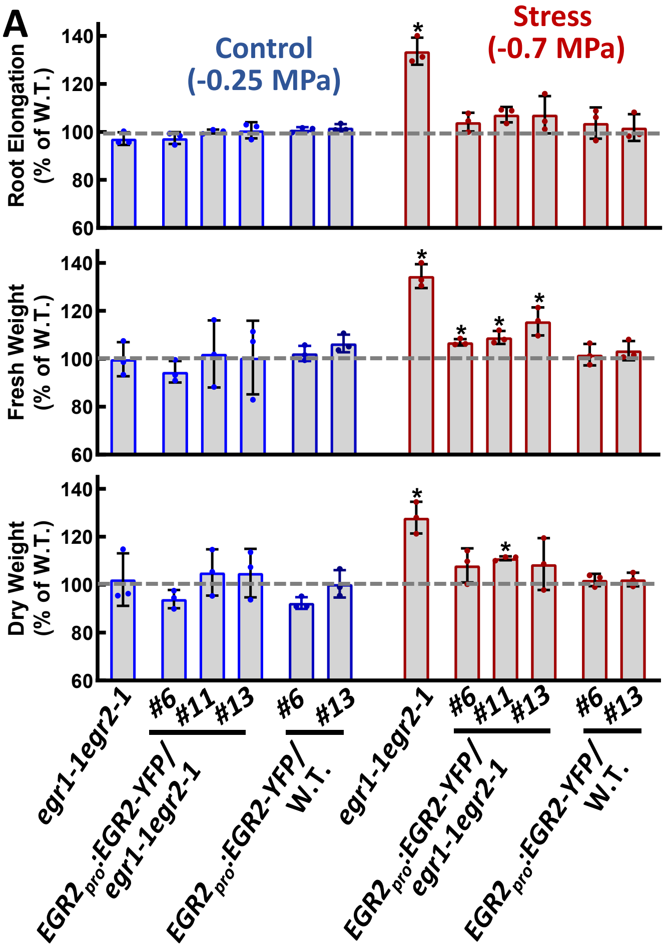


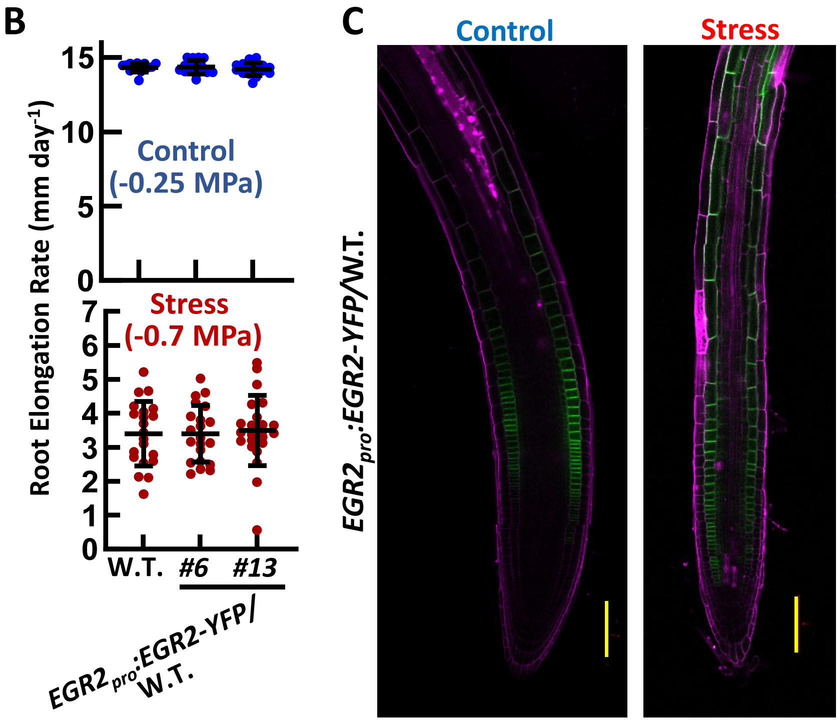


**Supplemental Figure S11: Complementation of *egr1-1egr2-1* by *EGR2$_{pro}$:EGR2-YFP* and EGR2 overexpression (*EGR2$_{pro}$:EGR2-YFP* in the wild type background) (Supports Fig. 4).**

**A.** Seedling primary root elongation, fresh weight and dry weight were assayed in unstressed control or low $\psi_w$ stress (-0.7 MPa). Asterisks indicate significant difference from wild type (100%, indicated by the dashed line) by one-sample T-test ($p \leq 0.05$). Data are combined from three independent experiments (n = 3). Note that the complementation lines still had higher than wild type levels of growth in some cases, presumably reflecting the effect of the *egr1-1* mutation.

**B.** Root elongation rates under control and stress treatments for W.T. and EGR2 overexpression (native promoter) transgenic lines. Data are combined from two biological experiments. Error bars indicate S.D. (n = No significant difference were detected.

C. Representative root tip images of the *EGR2$_{pro}$:EGR2-YFP/W.T.* transgenic lines shown in panel B. YFP fluorescence is shown in green, PI staining of cell wall is shown in magenta. Scale bars indicate 100 µm. For stress treatment five-day-old seedlings were transferred to -0.7 MPa PEG-agar plates and images 4 days after transfer. Seedlings of the same age of the same age were images for the unstressed control.

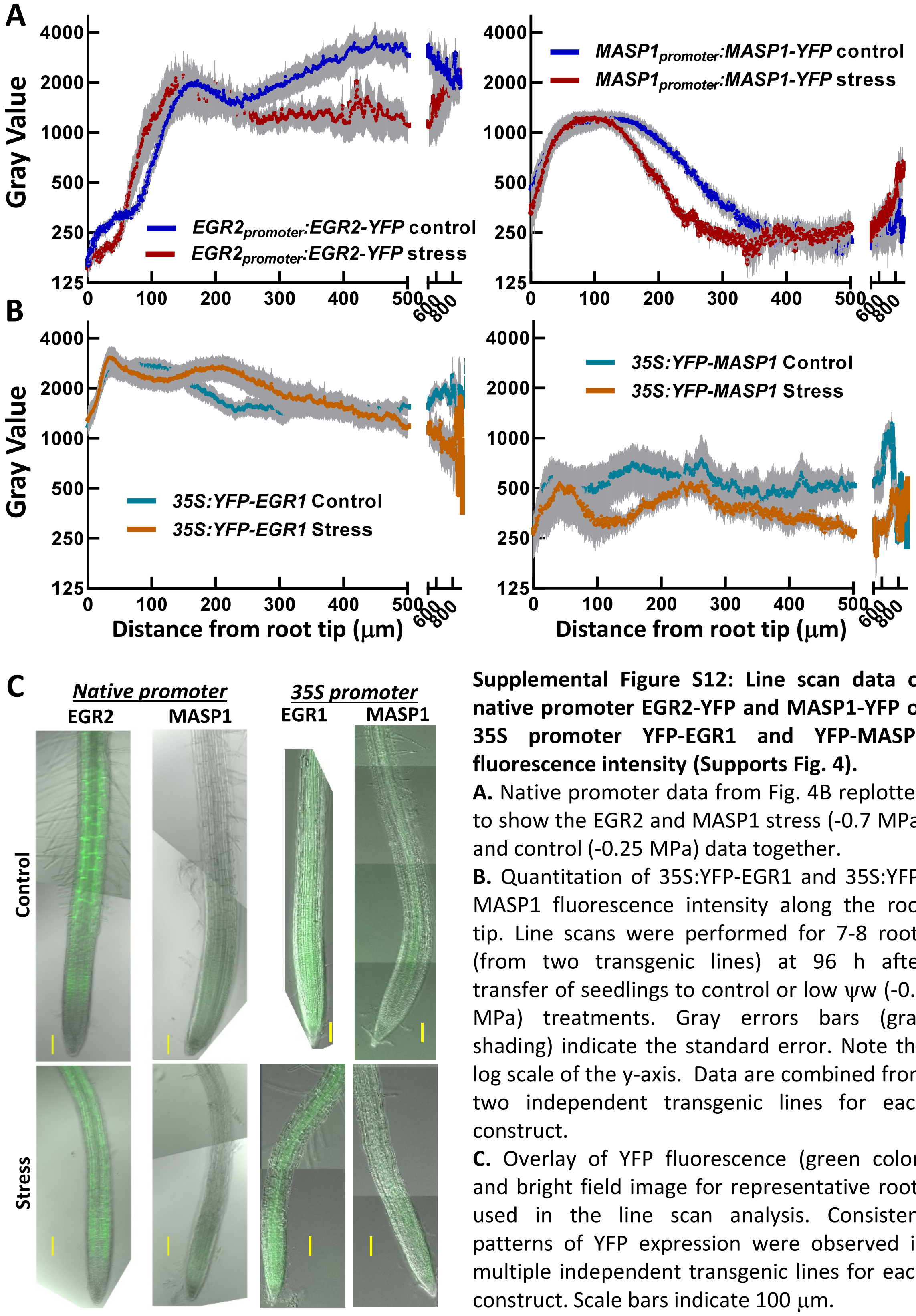


**Supplemental Figure S12: Line scan data of native promoter EGR2-YFP and MASP1-YFP or 35S promoter YFP-EGR1 and YFP-MASP1 fluorescence intensity (Supports Fig. 4).**

**A.** Native promoter data from Fig. 4B replotted to show the EGR2 and MASP1 stress (-0.7 MPa) and control (-0.25 MPa) data together.

**B.** Quantitation of 35S:YFP-EGR1 and 35S:YFP-MASP1 fluorescence intensity along the root tip. Line scans were performed for 7-8 roots (from two transgenic lines) at 96 h after transfer of seedlings to control or low ψw (-0.7 MPa) treatments. Gray errors bars (gray shading) indicate the standard error. Note the log scale of the y-axis. Data are combined from two independent transgenic lines for each construct.

**C.** Overlay of YFP fluorescence (green color) and bright field image for representative roots used in the line scan analysis. Consistent patterns of YFP expression were observed in multiple independent transgenic lines for each construct. Scale bars indicate 100 μm.

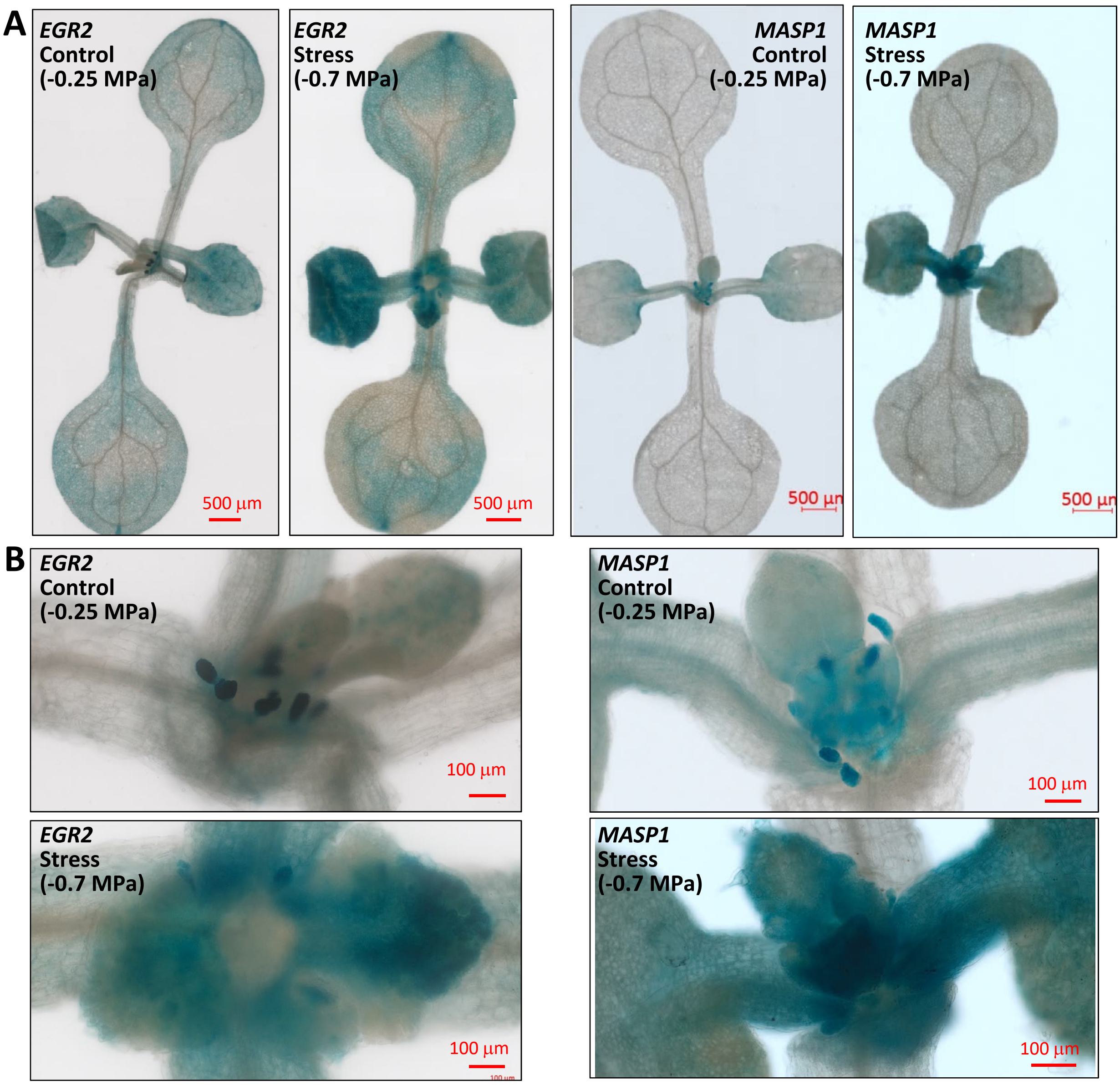


**Supplemental Figure S13: *EGR2* and *MASP1 promoter:GUS* expression pattern in shoot meristem and young leaf (Supports Fig. 5).**

A. Shoot GUS staining in 6 day old seedlings (control) or 9-day-old seedlings under stress conditions (4 days after transfer of 5-day-old seedlings to -0.7 MPa stress plates). The different ages were used so that seedlings of the same size could be imaged for both treatments. $EGR2_{promoter}$:*GUS* activity was detected, and was induced by low water potential stress in young leaves but not in the shoot meristem. $MASP1_{promter}$:*GUS* was detected around the shoot meristem and at the base of young leaves. Similar expression pattern was observed in multiple independent transgenic lines. Scale bars indicate 500 μm.

B. Close up of shoot meristem regions to better illustrate $EGR2_{promoter}$:*GUS* in leaf primordia and young leaf but not in shoot meristem while conversely $MASP1_{promter}$:*GUS* is most highly expressed in the meristem under both control and stress conditions. Scale bars indicate 100 μm.

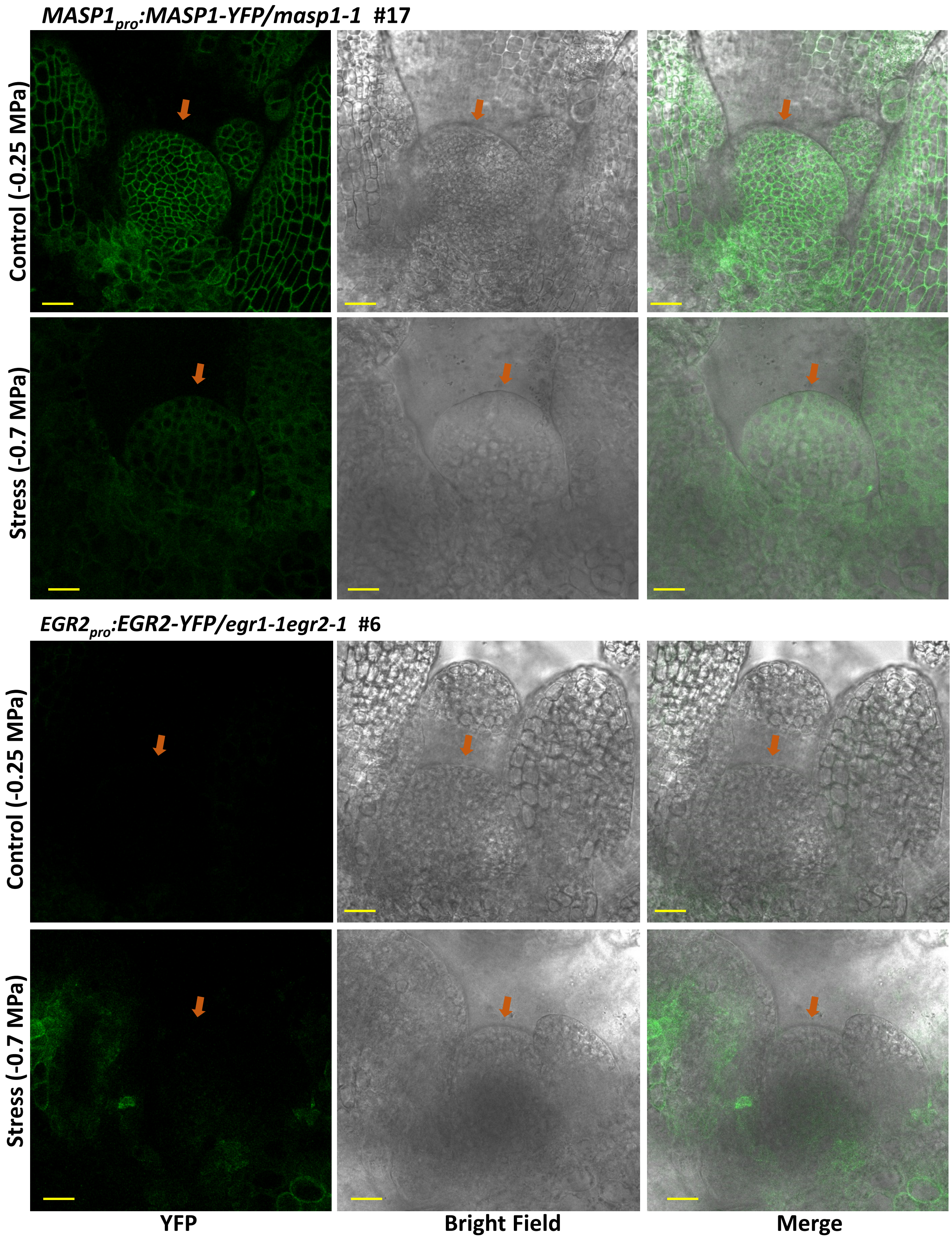


**Supplemental Figure S14 (legend on next page)**

**Supplemental Figure S14 (previous page): EGR2 and MASP1 expression in shoot meristem (Supports Fig. 5).**

Imaging was performed on 6 day old seedlings (control) or 9-day-old seedlings under stress conditions (4 days after transfer of 5-day-old seedlings to -0.7 MPa stress plates). The different ages were used so that seedlings of the same size could be imaged for both treatments. Scale bars indicate 20 $\mu$m. Orange arrows point to the top of the shoot meristem (surrounding tissues are mainly leaf primordia).

Expression of *EGR2pro:EGR2-YFP* was not detected in shoot meristem under either control or stress conditions. Conversely, *MASP1pro:MASP1-YFP* could be detected in shoot meristem in both treatments the MASP1-YFP signal tended to be weaker in the shoot meristem after low $\psi$w treatment.

Images shown are representative of the expression pattern observed in three independent transgenic lines for each construct.
